# Substrate-induced asymmetry coordinates lipoprotein handover and abaucin trapping in *Acinetobacter baumannii* LolDF

**DOI:** 10.64898/2026.10.30.762977

**Authors:** Jianping Kong, Ziyi Zhu, Zhi Lin, Nan Yao, Huang Ye, Hongfeng Gu, Ruiyan He, Meirong Chen, Meiling Lu, Haifeng Sun, Yibei Xiao, Zhaoxing Li

## Abstract

The homodimeric lipoprotein transporter LolDF supports outer-membrane biogenesis in *Acinetobacter baumannii*. How homodimeric LolDF generates directional lipoprotein transport, and how the reported two- and four-abaucin states are related, remain unclear. Using defined lipidated *Ab*LolB and *Ab*LolA, we show that substrate binding breaks LolF symmetry: LolF_B_ primarily accommodates the lipid anchor, LolF_A_ positions the adjacent polypeptide, and LolA docks asymmetrically through the LolF hook and an *Ab*LolA-specific loop. C1 reconstruction of the wide abaucin state reveals non-equivalent central ligands, whereas C2 processing of the same particles averages this asymmetry; the two peripheral ligands form a stable inhibitory core, while the central sites are conditionally occupied in the expanded cavity. Abaucin inhibits ATPase activity and LolA-mediated transfer from substrate-associated LolDF, supporting a unified mechanism in which substrate-induced asymmetry organizes productive handover and its conformational plasticity is redirected during inhibition.

## Introduction

Antimicrobial resistance is reducing treatment options for severe Gram-negative infections. In 2021, bacterial antimicrobial resistance was associated with 4.71 million deaths worldwide^1^. Carbapenem resistance rose especially quickly among Gram-negative pathogens^1^. The World Health Organization lists carbapenem-resistant *Acinetobacter baumannii* as a critical-priority pathogen^2^. The outer membrane helps these bacteria survive antibiotics and other environmental stresses^3^. Many proteins that build and maintain this membrane are lipoproteins anchored by an N-terminal lipid. Outer-membrane lipoproteins must leave the inner membrane and cross the periplasm to reach their destination. The essential transport machinery that drives this pathway offers targets for new antibacterial agents^4–6^.

The Lol pathway is widely distributed across Gram-negative bacteria and directs outer-membrane lipoproteins from the inner membrane to their final destination. Its ATP-binding cassette (ABC) transporter in the inner membrane occurs in two main forms, heterodimeric LolCDE and homodimeric LolDF^7^. The best-characterized system is *Escherichia coli* LolCDE. LolC and LolE form an asymmetric transmembrane heterodimer, while two LolD subunits provide cytoplasmic ATPase activity^8,9^. LolCDE captures lipidated cargo at the LolC-LolE interface and releases it to the periplasmic chaperone LolA. LolA then carries the lipoprotein to the outer-membrane receptor LolB^10,11^. Structures of cargo-, nucleotide- and LolA-bound LolCDE show that LolC and LolE make different contributions to transport^9,12–15^.

*A. baumannii* uses the homodimeric transporter LolDF, in which two copies of LolF replace the specialized LolC and LolE subunits of LolCDE. This architecture removes the fixed sequence asymmetry that underlies current models of LolCDE transport. Recent structures have begun to define the LolDF transport cycle, but key steps remain unresolved. Zhang *et al.* captured *A. baumannii* LolDF in nucleotide-free, vanadate-trapped and inhibitor-bound conformations, establishing a framework for its conformational cycle, but did not resolve a lipoprotein-bound or LolA-bound complex^16^. Pang *et al.* subsequently determined lipoprotein-like substrate-, AMP-PNP- and LolA-bound states of *A. baylyi* LolDF^17^. However, this work used a different *Acinetobacter* species, and the endogenous lipoprotein captured during purification could not be assigned unambiguously. Thus, how *A. baumannii* LolDF recognizes a defined lipidated substrate and organizes its transfer to LolA remains unclear.

LolDF has also emerged as a species-selective antibacterial target. Abaucin, identified by a machine-learning-guided screen, selectively inhibits *A. baumannii* lipoprotein trafficking^18^. Two recent structural studies captured wide- and narrow-cavity abaucin-bound states but reached different descriptions of inhibitor binding and action. Zhang *et al.* resolved two- and four-abaucin complexes and proposed that sequential binding drives stepwise, induced-fit opening of the transmembrane cavity^16^. Pang *et al.* instead modelled two symmetrically arranged abaucin molecules in both conformations and proposed that abaucin traps a substrate-accessible but transport-incompetent state that prevents transfer to LolA^19^. The wide-state ligand arrangements and resulting mechanistic models therefore differ, and the relationship between the wide and narrow states remains unresolved.

Here we address these gaps directly in *A. baumannii* LolDF. We determine a complex containing a defined lipidated *Ab*LolB substrate and show that a single lipoprotein breaks the apparent symmetry of the LolF homodimer, with the two identical protomers making distinct contributions to lipid-anchor and polypeptide recognition. *Ab*LolA-engaged structures and interface perturbations further define the docking geometry required for productive handover. We also recover four- and two-abaucin states and show, by processing the same wide-state particles under C1 and C2 symmetry, that non-equivalent central ligand configurations can be obscured by symmetry averaging. The peripheral inhibitor sites are retained as the cavity narrows, providing a structural connection between the previously reported two- and four-ligand states. Together with functional measurements showing inhibition of transfer from substrate-associated LolDF, these results link substrate-induced asymmetry to LolA-mediated handover and provide a framework for understanding how abaucin traps non-productive LolDF conformations.

## Results

### Nucleotide binding drives conformational rearrangements of *Ab*LolDF

We purified wild-type (WT) *Ab*LolDF and two LolD catalytic mutants, E172Q and H205A (Extended Data Fig. 1a). Size-exclusion chromatography and SDS-PAGE showed similar elution profiles and subunit patterns for all three complexes. Both mutants retained the overall subunit assembly. LC-MS/MS identified LolD and LolF as the dominant proteins in the SEC-purified WT sample. The same analysis detected multiple *Escherichia coli* lipoproteins at lower abundance (Extended Data Fig. 1b).

We measured the ATPase activity of WT *Ab*LolDF and both LolD variants. WT *Ab*LolDF showed clear ATP-dependent ATPase activity (Fig. 1b). The activity approached saturation as the ATP concentration increased. By contrast, the activities of E172Q and H205A remained close to background across the tested ATP range. E172 is the conserved catalytic glutamate adjacent to the Walker B motif. This glutamate participates in the catalytic water network^20^. H205 is the conserved H-loop, or switch, histidine. The H-loop histidine helps organize the catalytic site and contacts the ATP γ-phosphate^21^. The loss of ATPase activity supports the use of E172Q to capture a hydrolysis-deficient, ATP-bound state.

**Fig. 1.**
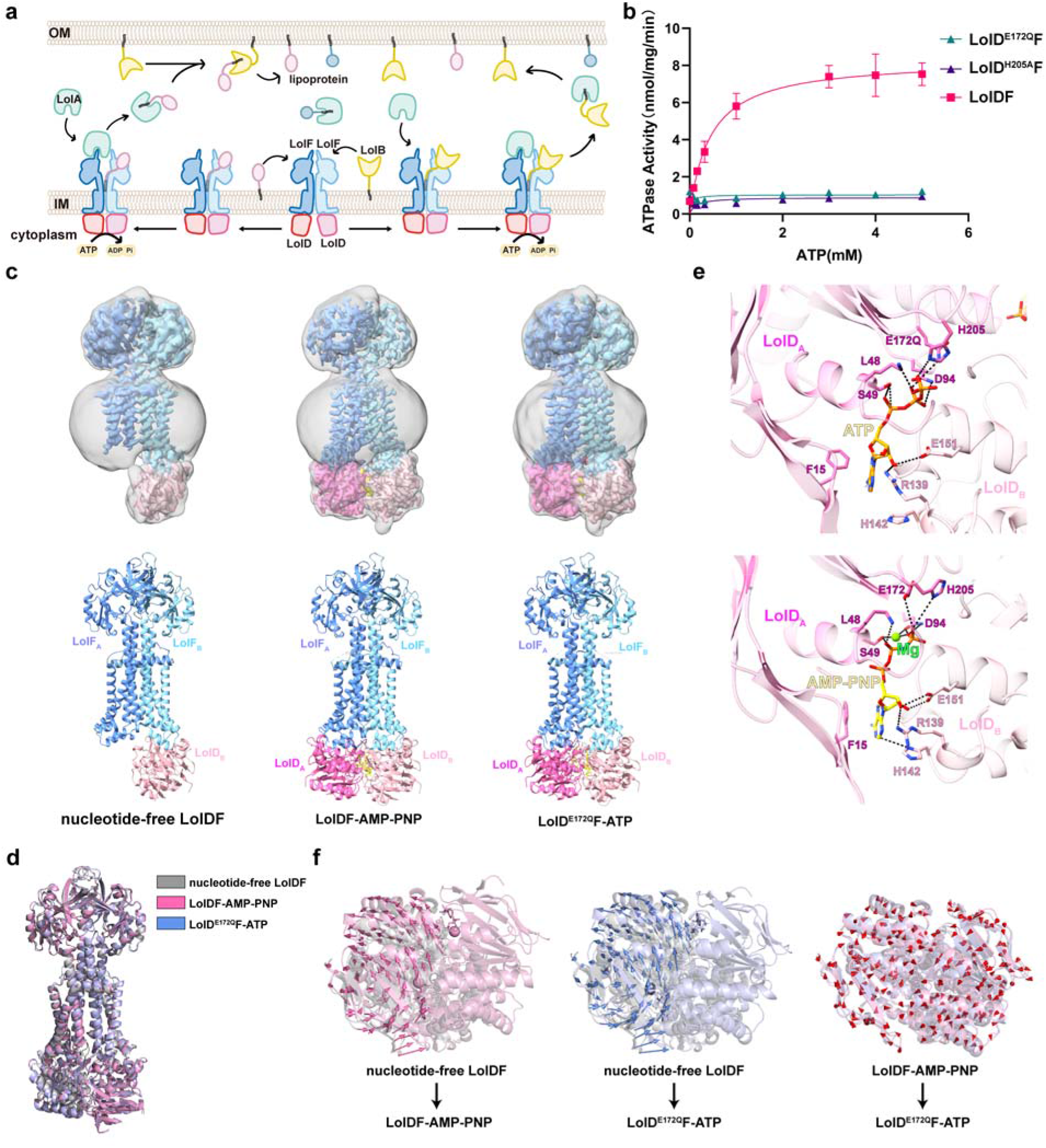
Nucleotide-dependent conformational states of *Ab*LolDF. **a,** Schematic of the Lol pathway showing lipoprotein extraction by LolDF, transfer to LolA and subsequent delivery to LolB. **b,** ATP-dependent ATPase activities of wild-type *Ab*LolDF and the *Ab*LolD^E172Q^F and *Ab*LolD^H205A^F variants. Data are presented as mean ± s.d. from three independent experiments, each performed in technical triplicate. **c,** Cryo-EM density maps (top) and atomic models (bottom) of nucleotide-free *Ab*LolDF, *Ab*LolDF-AMP-PNP and *Ab*LolD^E172Q^F-ATP. Grey densities represent Gaussian-smoothed maps displayed in UCSF ChimeraX (width = 2.0). **d,** Structural superposition of the LolD subunits in the three indicated conformational states. **e,** Close-up views of the ATP- and AMP-PNP-binding sites. Hydrogen bonds are shown as black dashed lines. **f,** Displacement-vector analysis of LolD between the indicated conformational states. Vector direction indicates the direction of local structural displacement, and vector length is proportional to the magnitude of movement.

We next determined cryo-EM structures of nucleotide-free *Ab*LolDF (3.4 Å), WT *Ab*LolDF-AMP-PNP (2.8 Å) and *Ab*LolD^E172Q^F-ATP (3.6 Å) (Fig. 1c, Extended Data Figs. 2 and 3). The three structures retain the same overall architecture, but the relative arrangement of the cytoplasmic LolD subunits differs among states (Fig. 1d). In both nucleotide-bound structures, two nucleotide molecules occupy composite sites at the LolD dimer interface (Fig. 1e). One LolD subunit contributes the Walker A and Walker B motifs, E172 and H205. The opposing LolD subunit completes the site through its ABC signature motif ^20,22^ (Extended Data Fig. 1c). The WT *Ab*LolDF-AMP-PNP map also contains density for Mg^2+^. The *Ab*LolD^E172Q^F-ATP map retains clear ATP density but lacks corresponding Mg^2+^ density. E172Q therefore retained ATP binding despite its loss of hydrolytic activity.

Structural superposition and displacement-vector analysis defined the conformational changes associated with nucleotide binding (Fig. 1d and Fig. 1f). The LolF transmembrane region retains a similar overall framework across the three structures. The largest movements occur within the cytoplasmic LolD assembly. Nucleotide binding brings the two LolD subunits closer and reorganizes their dimer interface around the composite nucleotide-binding sites. The *Ab*LolD^E172Q^F-ATP and WT *Ab*LolDF-AMP-PNP structures also adopt distinct conformations. The displacement vectors reveal coordinated motion across several LolD regions. LolD closure was accompanied by local rearrangements within the dimer. We next examined lipidated LolB recognition by ATP-bound *Ab*LolDF.

### *Ab*LolDF recognizes lipidated *Ab*LolB through a composite transmembrane cavity

We purified *Ab*LolDF in complex with lipidated *Ab*LolB and examined the complex by cryo-EM in the presence of ATP (Fig. 2a). Size-exclusion chromatography and SDS-PAGE showed that *Ab*LolDF and *Ab*LolB formed a stable complex (Extended Data Fig. 4a). Two-dimensional classification, heterogeneous refinement and final refinement yielded a reconstruction at an overall resolution of approximately 3.1 Å. The reconstruction contains two LolF subunits, two LolD subunits and density for one lipidated *Ab*LolB molecule. The lipidated N-terminal cysteine C22, the following 17 residues and the three acyl chains attached to C22 were resolved sufficiently for atomic modelling. The remainder of *Ab*LolB was visible as additional density in a lower-resolution map but was not included in the atomic model (Fig. 2a). This density places the globular domain of *Ab*LolB on the periplasmic side of the transporter. The modelled lipidated N terminus extends into the central cavity between the two LolF protomers. This ATP-bound complex captures *Ab*LolB engaged in the central cavity.

**Fig. 2.**
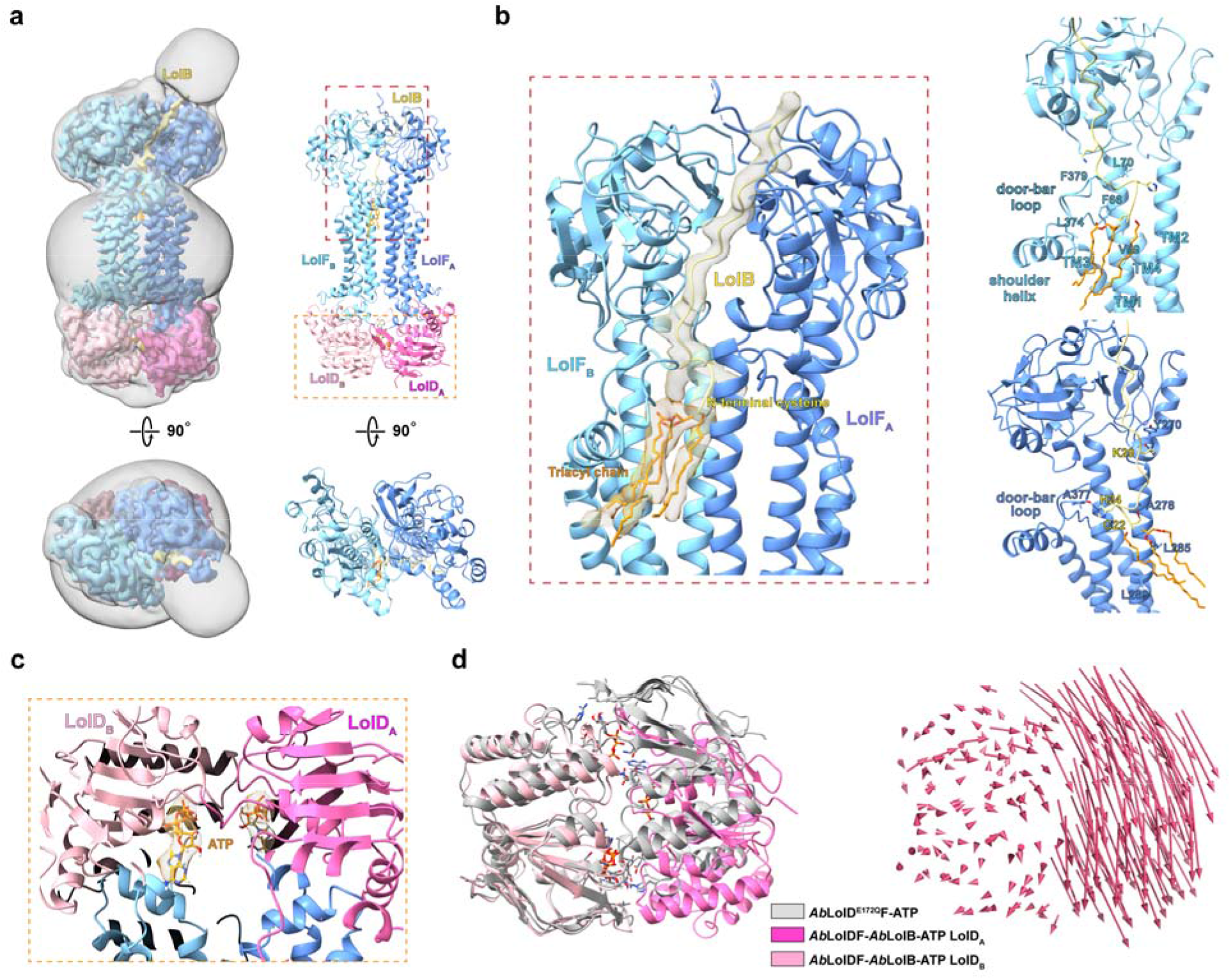
Structural basis for recognition of lipidated *Ab*LolB by *Ab*LolDF. **a,** Cryo-EM density map (left) and atomic model (right) of the ATP-bound *Ab*LolDF-*Ab*LolB complex, viewed from the membrane plane and the periplasm. Grey density represents a Gaussian-smoothed map displayed in UCSF ChimeraX (width = 2.0). **b,** Close-up view of lipidated *Ab*LolB within the central cavity of *Ab*LolDF. The right panels show representative interactions between the *Ab*LolB N terminus and the two LolF protomers. **c,** Close-up view of an ATP-binding site in the *Ab*LolDF-*Ab*LolB complex. **d,** Structural superposition and displacement-vector analysis of the LolD subunits in *Ab*LolDF-*Ab*LolB-ATP and *Ab*LolD^E172Q^F-ATP. Vector direction indicates the direction of local displacement, and vector length is proportional to the magnitude of movement.

The two LolF protomers jointly recognize the lipid anchor and the N-terminal polypeptide of *Ab*LolB, but they make different contributions to substrate binding (Fig. 2b). The mature N terminus of *Ab*LolB enters the central cavity through the interface between the two LolF subunits. The lipidated C22 residue and its three acyl chains are buried within the transmembrane region. This lipidated segment lies closer to the door-bar loop of LolF_B_ than to the corresponding loop of LolF_A_. LolF_B_ forms a larger set of hydrophobic contacts with the lipidated N terminus. V59, F66, L70, L374 and F379 from LolF_B_ line this side of the cavity and surround the acyl chains. By contrast, LolF_A_ interacts more closely with the N-terminal polypeptide of *Ab*LolB. LolF_A_ contributes Y270, A278, L285, L289 and A377 to the interface. The backbone carbonyl oxygen of Y270 accepts a hydrogen bond from the side-chain nitrogen of *Ab*LolB K29. The side-chain nitrogen of *Ab*LolB H24 forms a second hydrogen bond with the backbone of A377. The remaining residues line the cavity and provide additional hydrophobic contacts. LolF_B_ primarily accommodates the lipid anchor, while LolF_A_ positions the adjacent polypeptide segment. We tested this composite interface using a panel of structure-guided LolF mutants. The *Ab*LolDF variants could be purified, but none of them co-purified with the lipoprotein cargo (Extended Data Fig. 5a).

Both nucleotide-binding sites contained clear density for intact ATP, including the γ-phosphate (Fig. 2c). Each ATP molecule is clamped between the two LolD subunits. One LolD contributes the Walker A and Walker B motifs, while the opposing LolD contributes the ABC signature motif^22^. The two ATP molecules stabilize a compact LolD dimer in the substrate-engaged complex. Although the complex was incubated with 2 mM ATP and 1 mM MgCl_2_ during cryo-EM sample preparation, no well-defined Mg^2+^ density was resolved at either catalytic site.

The two LolD monomers superimposed closely despite one-sided *Ab*LolB engagement (Extended Data Fig. 5b). We next compared the LolD dimer in the ATP-bound *Ab*LolDF-*Ab*LolB complex with that in *Ab*LolD^E172Q^F-ATP (Fig. 2d and Extended Data Fig. 5c). Both structures contained two ATP molecules, but the relative arrangement of the LolD pair differed between the two states. Alignment of one LolD subunit revealed a coordinated displacement of the opposing subunit. The displacement vectors also identified local movements across several regions of the LolD dimer. The ATP-bound complexes therefore differed in LolD dimer arrangement.

The two LolF protomers also showed different degrees of opening in the *Ab*LolB-engaged structure (Extended Data Fig. 5d). Comparison with *Ab*LolD^E172Q^F-ATP revealed local shifts in the transmembrane helices and door-bar loops surrounding the central cavity. The door-bar loop of LolF_B_ showed a larger displacement than the corresponding loop of LolF_A_. This loop lies next to the N-terminal acyl chains of *Ab*LolB. Its displacement widens the LolF_B_ side of the cavity and provides an opening for the lipidated N terminus to enter. LolF_A_ shows a smaller opening and forms more direct contacts with the polypeptide segment. These structural differences create an asymmetric substrate-binding cavity within the LolF homodimer.

### LolA engages the substrate-loaded *Ab*LolDF complex in two conformational states

We next examined *Ab*LolA docking onto substrate-loaded *Ab*LolDF. We assembled *Ab*LolDF-*Ab*LolB with GST-tagged *Ab*LolA and purified the complex by size-exclusion chromatography. SEC and SDS-PAGE showed that the proteins formed a stable complex (Extended Data Fig. 6a). Three-dimensional classification resolved two conformational states (Extended Data Fig. 6b-h). State 1 was reconstructed at an overall resolution of approximately 2.9 Å (Fig. 3a), whereas state 2 was reconstructed at approximately 2.7 Å (Fig. 3b). Both states contain *Ab*LolDF, the lipidated N-terminal region of *Ab*LolB and *Ab*LolA (Extended Data Fig. 7a). *Ab*LolB remains at the periplasmic entrance of the central cavity, with *Ab*LolA positioned adjacent to its lipidated N terminus. The two structures share a similar overall architecture but differ more substantially in the cytoplasmic LolD assembly.

**Fig. 3.**
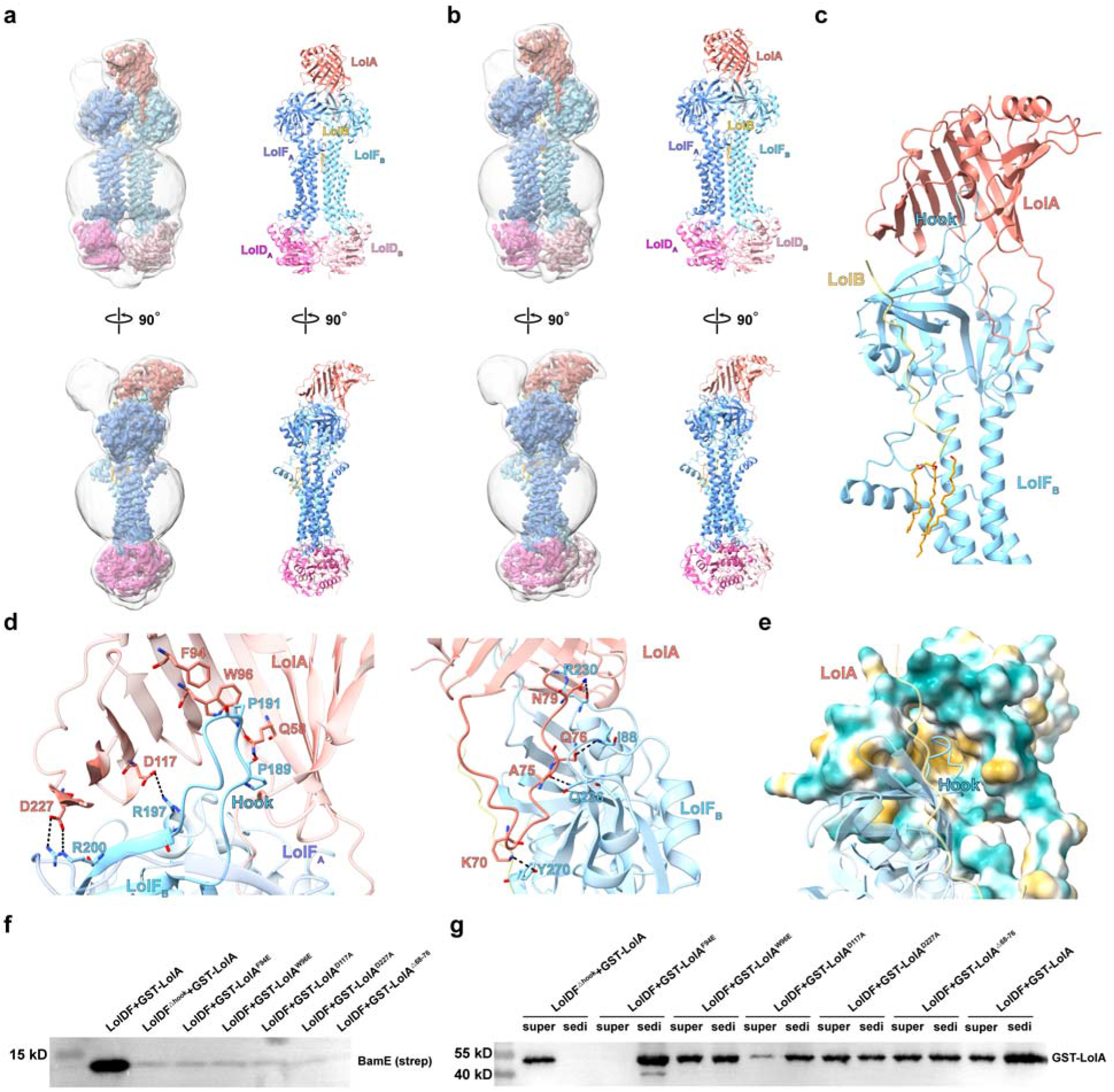
Cryo-EM structures of *Ab*LolDF-*Ab*LolB-GST-*Ab*LolA in two conformational states. **a,** Cryo-EM density map (left) and atomic model (right) of state 1. Grey density represents a Gaussian-smoothed map displayed in UCSF ChimeraX (width = 2.0). **b,** Cryo-EM density map (left) and atomic model (right) of state 2. Grey density represents a Gaussian-smoothed map displayed in UCSF ChimeraX (width = 2.0). **c,** Overall view of the interfaces formed by LolF protomer B with *Ab*LolA and *Ab*LolB. **d,** Close-up views of polar interactions between *Ab*LolA and the two LolF protomers. Hydrogen bonds are shown as black dashed lines. The *Ab*LolA-specific loop spans residues 68-79; the Δ68-76 variant removes part of this loop. **e,** Cross-sectional hydrophobicity surface of the *Ab*LolA-*Ab*LolDF interface. Hydrophilic and hydrophobic surfaces are coloured turquoise and goldenrod, respectively. **f,** Representative immunoblot showing in vitro lipoprotein transfer from *Ab*LolDF-BamE proteoliposomes to wild-type or the indicated *Ab*LolA interface variants. A representative result from three independent experiments is shown; each experiment was performed in technical triplicate, with similar results. **g,** Proteoliposome co-sedimentation analysis of wild-type and the indicated *Ab*LolA variants with *Ab*LolDF. A representative immunoblot from three independent experiments is shown. Super, supernatant; sedi, sediment.

The periplasmic regions of the two LolA-engaged states superimpose closely. We therefore used one state as a representative structure to analyse the intermolecular contacts. *Ab*LolA docks across a composite platform formed by the periplasmic domains of both LolF protomers (Fig. 3c). The two LolF protomers contribute non-equivalent interfaces. In LolF_B_, residues 186-196 form a periplasmic hook that projects into the hydrophobic cavity of *Ab*LolA (Fig. 3d). F94 and W96 of *Ab*LolA pack against the hook near P191 of LolF_B_, while Q58 forms a hydrogen bond with the backbone of P189. Additional polar interactions connect D117 and D227 of *Ab*LolA with R197 and R200 of LolF_B_, respectively. These contacts anchor the LolF_B_ hook within the *Ab*LolA cavity. LolF_A_ forms a smaller secondary interface with a separate surface of *Ab*LolA (Extended Data Fig. 7c).

A second interface is formed by an *Ab*LolA-specific loop spanning residues 68-79 (Fig. 3d). This loop lies against the periplasmic domain of LolF_B_, with contacts between K70 of *Ab*LolA and Y270 of LolF_B_, A75 and Q236, Q76 and I88, and N79 and R230. The hook-binding site and the 68-79 loop therefore engage distinct regions of LolF_B_ and together position *Ab*LolA above the substrate-binding cavity. The hydrophobic cavity of *Ab*LolA faces the lipidated N terminus of *Ab*LolB and the central cavity of *Ab*LolDF (Fig. 3e), creating a handover geometry between the transporter-bound substrate and the LolA cavity. The acyl chains remain associated with *Ab*LolDF in both structures, with the *Ab*LolA cavity positioned to receive them.

To test the functional contribution of the *Ab*LolA-specific loop, we generated an *Ab*LolA Δ68-76 variant, which removes most of the segment contacting the LolF periplasmic domain, including K70, A75 and Q76. Deletion of residues 68-76 strongly reduced lipoprotein transfer to *Ab*LolA (Fig. 3f), but the variant retained detectable association with *Ab*LolDF in the proteoliposome co-sedimentation assay (Fig. 3g). The loop contributes to productive handover without being essential for detectable *Ab*LolA association.

We next tested the transporter-side docking element. Deletion of the LolF hook spanning residues 186-196 nearly abolished lipoprotein transfer in the reconstituted system (Fig. 3f), while the truncated *Ab*LolDF complex retained substantial ATPase activity (Extended Data Fig. 7e). The LolF hook therefore couples ATP turnover to productive LolA engagement and substrate handover.

Point mutations within the *Ab*LolA docking surface produced distinct phenotypes. W96E, D117A and D227A reduced association with *Ab*LolDF and strongly impaired lipoprotein transfer (Fig. 3f, g), consistent with disruption of the structurally observed interface. F94E produced the opposite binding phenotype. In the proteoliposome co-sedimentation assay, most WT *Ab*LolA remained in the soluble fraction after incubation and ultracentrifugation, whereas *Ab*LolA^F94E^ predominantly co-sedimented with *Ab*LolDF proteoliposomes (Fig. 3g). Despite this persistent association, *Ab*LolA^F94E^ failed to support efficient lipoprotein transfer (Fig. 3f). These variants link the organization and persistence of LolA docking to productive lipoprotein transfer.

### Conformational changes associated with lipoprotein transport by *Ab*LolDF

We compared five *Ab*LolDF structures comprising nucleotide-free *Ab*LolDF, *Ab*LolDF-AMP-PNP, *Ab*LolDF-*Ab*LolB-ATP and the two *Ab*LolA-engaged states (Fig. 4a). Nucleotide binding was associated with rearrangement of the cytoplasmic LolD dimer. *Ab*LolB engagement was associated with a larger change that extended into the LolF transmembrane region. Both *Ab*LolA-bound states retained the open periplasmic architecture but differed at the cytoplasmic coupling interface. Measurements across the LolD dimer further showed that its upper and lower regions moved to different extents (Extended Data Fig. 8a). The two LolD subunits therefore undergo coupled closure and rotation rather than a uniform rigid-body movement.

**Fig. 4.**
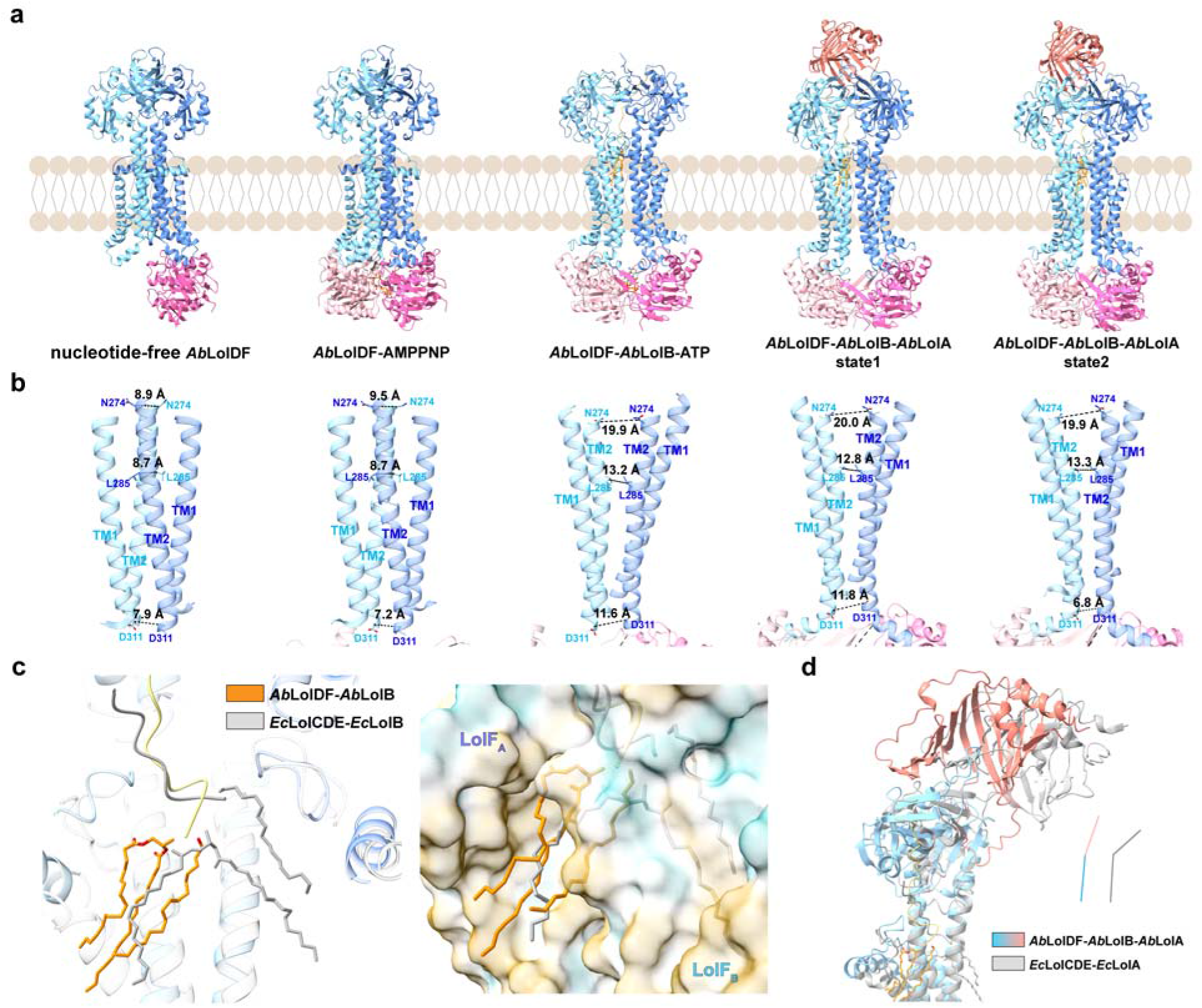
Conformational changes associated with lipoprotein transport by *Ab*LolDF. **a,** Cryo-EM structures of *Ab*LolDF in the nucleotide-free, AMP-PNP-bound, *Ab*LolB-bound and two *Ab*LolA-bound states. LolF protomers A and B are coloured cornflower blue and light sky blue, respectively. LolD protomers A and B are coloured hot pink and light pink, respectively. *Ab*LolA is coloured salmon and *Ab*LolB is coloured khaki. **b,** Comparison of TM1 and TM2 across the indicated states. Distances between representative residues in the upper, middle and lower regions quantify changes in the transmembrane cavity. Distances were measured between corresponding Cα atoms of the two LolF protomers. **c,** Comparison of the triacylated N-terminal cysteine of *Ab*LolB bound to *Ab*LolDF with that of LolB bound to *Escherichia coli* LolCDE (PDB 9RLH; LolB Δ9-15). A cross-sectional hydrophobicity surface of the *Ab*LolDF cavity is shown on the right. **d,** Comparison of *Ab*LolDF-*Ab*LolB-*Ab*LolA with the *Escherichia coli* LolCDE-lipoprotein-LolA complex (PDB 7ARM).

We quantified the changes in central cavity width. TM1 and TM2 from the two LolF protomers form the opposing walls of the central substrate-binding cavity. We measured Cα-Cα distances between equivalent residues on the two TM2 helices at three positions (Fig. 4b). The upper N274 pair reports the width of the periplasmic entrance. The middle L285 pair lies next to the distal ends of the *Ab*LolB acyl chains and reports the width of the lipid-accommodating region. The lower D311 pair reports structural changes near the LolD-LolF coupling interface. The N274 distance increased from 8.9 Å in nucleotide-free *Ab*LolDF and 9.5 Å in *Ab*LolDF-AMP-PNP to 19.9 Å in the *Ab*LolB-bound structure. This distance remained close to 20 Å in both *Ab*LolA-engaged states. The L285 distance increased from 8.7 Å to 13.2 Å after *Ab*LolB engagement and remained between 12.8 and 13.3 Å after *Ab*LolA binding. The lower D311 distance followed a different pattern. It increased from 7.2-7.9 Å in the nucleotide-free and AMP-PNP-bound structures to 11.6 Å in the *Ab*LolB-bound structure and 11.8 Å in *Ab*LolA-engaged state 1. It measured 6.8 Å in state 2. The central cavity therefore undergoes regional reshaping rather than uniform expansion or contraction.

We next compared the substrate-binding modes of *Ab*LolDF and *Ec*LolCDE. The triacylated anchors occupied markedly different positions in the two transporters (Fig. 4c). In *Ab*LolDF, all three *Ab*LolB acyl chains entered the same side of the central cavity. The hydrophobic surface facing LolF_B_ formed the main lipid-binding environment (Extended Data Fig. 8c). In *Ec*LolCDE, the three *Ec*LolB acyl chains were divided between the two sides of the cavity. One chain occupied the LolC-facing side, whereas the other two extended towards the LolE-facing side. The superposition shows the different acyl-chain distributions (Extended Data Fig. 8d). *Ab*LolDF therefore breaks the twofold symmetry of its homodimer after substrate binding. LolF_B_ provides the main environment for the lipid anchor, whereas LolF_A_ makes stronger contacts with the *Ab*LolB polypeptide. *Ec*LolCDE instead uses the built-in asymmetry of LolC and LolE to distribute the three acyl chains across both sides of its cavity. The globular domain of *Ab*LolB also occupies a different position. *Ab*LolB sits diagonally above the LolF dimer, whereas *Ec*LolB binds more laterally to *Ec*LolCDE (Extended Data Fig. 8b)^14^. These different orientations align with the distinct trajectories of the lipidated N termini and their acyl chains.

The two transporters also recruit LolA through different docking geometries. Structural superposition showed that *Ab*LolA and *Ec*LolA occupy different positions relative to their transporters (Fig. 4d)^9^. Their hydrophobic cavities therefore approach the lipoprotein-binding site from different angles. *Ab*LolA contains an additional loop spanning residues 68-79. This *Ab*LolA-specific loop contacts the periplasmic domain of LolF_B_ through K70, A75, Q76 and N79 (Fig. 3d), whereas *Ec*LolA lacks the corresponding insertion. The *Ab*LolA-specific loop is separate from the LolF periplasmic hook spanning residues 186-196, which enters the *Ab*LolA hydrophobic cavity. Deletion of *Ab*LolA residues 68-76 strongly impaired lipoprotein transfer despite retaining detectable association with *Ab*LolDF (Fig. 3f, g), supporting a functional role for this insertion in establishing the *Ab*LolDF-specific handover geometry. The LolF hook and the *Ab*LolA loop form separate contacts that support the observed docking geometry.

### Abaucin conformationally traps *Ab*LolDF and blocks lipoprotein transport

We next examined how abaucin alters the conformational cycle of *Ab*LolDF. We first incubated *Ab*LolDF that had not been supplemented with a defined lipoprotein with abaucin and ATP. Three-dimensional classification resolved wide- and narrow-cavity conformations, with the wide state forming the major particle population (Fig. 5a and Extended Data Fig. 9). The wide state was reconstructed at 3.6 Å resolution. By contrast, the *Ab*LolDF-*Ab*LolB sample yielded only a narrow-cavity state after incubation with abaucin and ATP. This structure was reconstructed at 3.4 Å resolution. The wide cavity accommodated four abaucin molecules, whereas the narrow cavity contained two molecules (Fig. 5b). The *Ab*LolDF preparation used without a defined lipoprotein was not necessarily substrate-free, as LC-MS/MS detected several low-abundance endogenous *E. coli* lipoproteins that co-purified with *Ab*LolDF (Extended Data Fig. 1b). By contrast, the *Ab*LolDF-*Ab*LolB sample contained a defined lipoprotein cargo, and Gaussian smoothing of the narrow-cavity reconstruction revealed weak putative lipoprotein-associated density above the central cavity (Extended Data Fig. 10a). The two preparations differed in the abaucin-bound states recovered by classification.

**Fig. 5.**
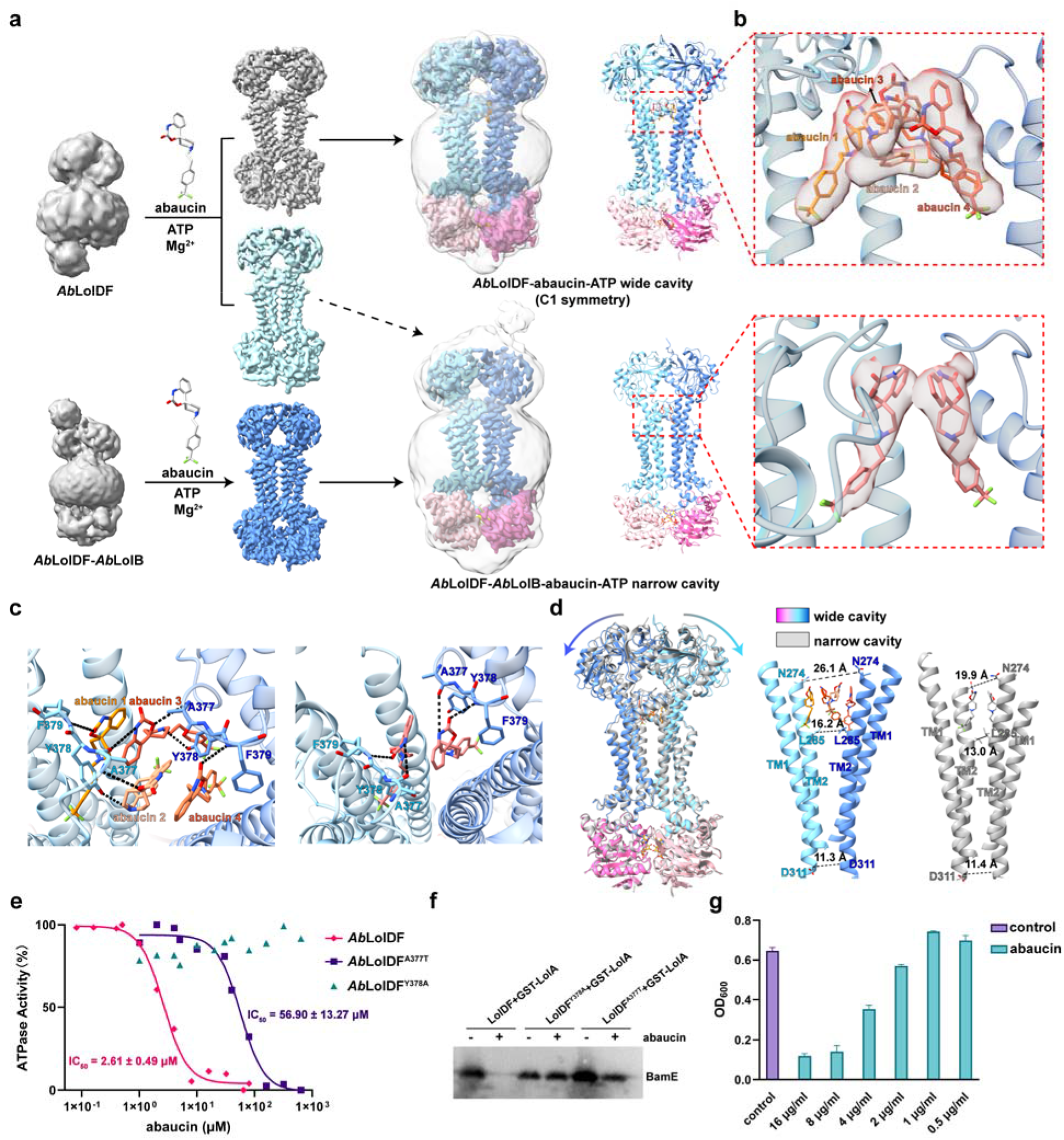
Structural and functional inhibition of *Ab*LolDF by abaucin. **a,** Cryo-EM maps and atomic models obtained from *Ab*LolDF-abaucin-ATP and *Ab*LolDF- *Ab*LolB-abaucin-ATP samples. *Ab*LolDF-abaucin predominantly adopts a wide-cavity state, whereas the *Ab*LolDF-*Ab*LolB-abaucin sample yields a narrow-cavity state. Grey densities represent Gaussian-smoothed maps displayed in UCSF ChimeraX (width = 2.0). **b,** Close-up views of abaucin densities in the wide- and narrow-cavity states. **c,** *Ab*aucin-binding interactions in the wide- and narrow-cavity states. Hydrogen bonds are shown as black dashed lines. **d,** Structural comparison of the wide- and narrow-cavity states. The right panels show TM1 and TM2, with representative Cα-Cα distances across the transmembrane cavity. **e,** *Ab*aucin concentration-response curves for the ATPase activities of wild-type *Ab*LolDF and the A377T and Y378A variants. Activities were normalized to the corresponding activities measured in the absence of abaucin. Data are presented as mean ± s.d. from three independent experiments, each performed in technical triplicate. Curves were fitted by nonlinear regression in Prism 10 (GraphPad). IC50 values are reported as best-fit values ± s.e. returned by the nonlinear regression; values are shown where a reliable fit could be obtained. **f,** Representative immunoblot showing abaucin inhibition of lipoprotein transfer from proteoliposomes reconstituted with preassembled *Ab*LolDF– BamE complexes to *Ab*LolA. A representative result from three independent experiments is shown; each experiment was performed in technical triplicate, with similar results. **g,** Growth of *Acinetobacter baumannii* ATCC 19606 after treatment with the indicated concentrations of abaucin for 4 h. Data are presented as mean ± s.d. from three biological replicates, each measured in technical triplicate.

The four abaucin molecules in the wide cavity did not adopt a strictly symmetric arrangement. The two peripheral molecules, abaucin 1 and abaucin 4, occupied binding sites and adopted poses closely resembling those of the two abaucin molecules in the narrow-cavity structure (Fig. 5d). These shared binding modes define two sites present in both the wide and narrow structures, whereas the two additional central molecules, abaucin 2 and abaucin 3, are accommodated only in the expanded wide cavity. The C1 reconstruction revealed distinct density patterns at these two central sites. The density contained both upright and bent components, consistent with at least two binding modes. Possible configurations include two extended ligands or a folded ligand, with an angle below 90°, paired with an extended ligand. To assess the effect of symmetry enforcement on interpretation of these densities, we reprocessed the same wide-state particle set with C2 symmetry as a processing control. C2 enforcement made the two central sites equivalent and produced similar intermediate density features at the two positions (Extended Data Fig. 10b,c). Thus, the C1 map retained non-equivalent ligand-density features that were averaged upon C2 symmetry enforcement. LolF residues A377, Y378 and F379 primarily contact the benzoxazine and piperidine moieties of abaucin (Fig. 5c), thereby anchoring the upper ring-containing core. By contrast, the trifluoromethyl phenyl group forms fewer contacts with LolF and is consequently less constrained. Consistent with this local flexibility, the density for the trifluoromethyl-containing end of abaucin 3 is comparatively weak and does not fully define this portion of the ligand. We propose that this weakly anchored end allows folded and extended poses to coexist at the two central sites. The narrow structure retains the two peripheral sites but lacks the additional central sites.

We next compared the abaucin-bound structures with the five uninhibited structures (Fig. 5d). We used the same N274, L285 and D311 distance measurements. The wide state showed the largest opening among all structures. Its upper N274 distance reached 26.1 Å, compared with 8.9 Å in nucleotide-free *Ab*LolDF, 9.5 Å in *Ab*LolDF-AMP-PNP and 19.9-20.0 Å in the substrate- and LolA-engaged states (Fig. 4b). Its middle L285 distance reached 16.2 Å, whereas the corresponding distance was 8.7 Å in the nucleotide-free and AMP-PNP-bound structures and 12.8-13.3 Å in the *Ab*LolB- and *Ab*LolA-bound structures. By contrast, its lower D311 distance was 11.3 Å. This value was close to those of *Ab*LolDF-*Ab*LolB-ATP and *Ab*LolA-engaged state 1, which measured 11.6 Å and 11.8 Å, respectively. The wide state therefore represents a selective expansion of the periplasmic entrance and lipid-accommodating region rather than uniform opening of the entire cavity.

The narrow state showed a different relationship with the transport cycle. Its N274, L285 and D311 distances were 19.9 Å, 13.0 Å and 11.4 Å, respectively. These values closely matched those of *Ab*LolDF-*Ab*LolB-ATP, which measured 19.9 Å, 13.2 Å and 11.6 Å. They also resembled *Ab*LolA-engaged state 1, which measured 20.0 Å, 12.8 Å and 11.8 Å. The narrow state matched *Ab*LolA-engaged state 2 at the upper and middle positions, but it lacked the strong lower-cavity contraction of state 2, where the D311 distance decreased to 6.8 Å. The narrow state therefore resembles the *Ab*LolB-bound and LolA-engaged state 1 structures at these TM2 positions, while differing from state 2 at D311.

Abaucin also produced a local change that was not captured by the three TM2 measurements. The distance between the opposing A377 positions increased from 14.5 Å in uninhibited *Ab*LolDF-*Ab*LolB-ATP to 18.1 Å in the abaucin-bound narrow state (Extended Data Fig. 10d). The retained abaucin molecules occupy a distinct arrangement around the door-bar loops, with greater separation at A377. The narrow state has substrate-like global cavity dimensions but retains an inhibitor-specific local geometry around the door-bar region. Its D311 separation also differs from that of LolA-engaged state 2.

Functional assays tested the role of the structurally defined binding site. Each ATPase dose-response curve was normalized to the corresponding activity in the absence of abaucin. Abaucin inhibited WT *Ab*LolDF with an IC_50_ of 2.61 ± 0.49 μM (Fig. 5e). The A377T substitution shifted the IC_50_ to 56.90 ± 13.27 μM, corresponding to an approximately 22-fold loss of sensitivity, whereas the Y378A variant retained ATPase activity across the tested abaucin range and did not yield a measurable IC_50_. The same substitutions protected lipoprotein transport from inhibition. Abaucin strongly reduced transfer mediated by WT *Ab*LolDF, whereas A377T and Y378A remained largely refractory to the compound (Fig. 5f). Abaucin also inhibited lipoprotein transfer from proteoliposomes reconstituted with preassembled *Ab*LolDF-BamE complexes, demonstrating that inhibition does not require prevention of initial substrate association. SPR measurements showed that abaucin bound directly to both *Ab*LolDF and *Ab*LolDF-*Ab*LolB, with similar apparent dissociation constants of 36.56 and 35.40 μM, respectively, across three independent experiments (Extended Data Fig. 10e). Abaucin also inhibited the growth of *A. baumannii* in a concentration-dependent manner (Fig. 5g).

The combined structural and functional data support two related routes of conformational trapping (Extended Data Fig. 10f). When lipoprotein occupancy is low, four abaucin molecules stabilize a wide cavity whose open periplasmic entrance may remain accessible to initial lipoprotein engagement. Complete insertion of the triacylated anchor would, however, probably require rearrangement or displacement of the two additional central molecules, abaucin 2 and abaucin 3, while preserving the two conserved binding modes represented by abaucin 1 and abaucin 4. Increased lipoprotein occupancy may therefore shift the conformational ensemble towards a contracted state in which only the two conserved abaucin molecules remain trapped. Importantly, abaucin also inhibited transfer from proteoliposomes reconstituted with preassembled *Ab*LolDF-BamE complexes (Fig. 5f). Because BamE was already associated with LolDF before abaucin exposure, inhibition does not require prevention of initial substrate association. These findings are consistent with abaucin engaging a substrate-associated transporter and stabilizing a non-productive state that cannot complete transfer to LolA. Thus, the wide and narrow structures may represent different routes into a common inhibited ensemble: abaucin can trap LolDF under different substrate-occupancy conditions, but in both cases uncouples substrate association from subsequent handover to LolA. Although the precise temporal relationship between the four-molecule wide state and the two-molecule narrow state remains to be established, both conformations divert *Ab*LolDF from its normal transport cycle.

## Discussion

Our study defines how the homodimeric *Ab*LolDF transporter coordinates lipoprotein recognition, LolA recruitment and inhibitor action. Previous mechanistic models of lipoprotein transport have largely been built around *E. coli* LolCDE, in which the divergent LolC and LolE subunits carry preassigned roles in substrate transport^9,14,15,23^. Previous *A. baylyi* structures resolved substrate-, nucleotide- and LolA-bound LolDF, including asymmetric substrate contacts within the homodimer^17^. Our structures sample a complementary set of states in *A. baumannii*, including a complex with defined lipidated *Ab*LolB and two *Ab*LolA-engaged conformations (Figs. 2a,b and 3a,b). Heterologously expressed *Ab*LolDF also co-purified with multiple *E. coli* lipoproteins (Extended Data Fig. 1b). Binding of a single defined lipoprotein breaks the apparent symmetry of the LolF homodimer, allowing two sequence-identical protomers to acquire distinct roles during substrate recognition and LolA engagement (Fig. 2b and Extended Data Fig. 5d; Fig. 3c-e and Extended Data Fig. 7c). LolCDE encodes different subunit roles through sequence divergence; in *Ab*LolDF, substrate contacts assign different roles to sequence-identical LolF protomers.

The conformational series suggests that nucleotide binding primarily reorganizes the cytoplasmic LolD motor (Fig. 1c-f), whereas substrate engagement produces the major opening and asymmetry of the LolF cavity (Figs. 2a,b and 4a,b; Extended Data Fig. 5c,d). Once *Ab*LolA docks, the periplasmic substrate-handover platform remains comparatively stable while additional rearrangements occur in the cytoplasmic and lower transmembrane regions (Fig. 3a,b; Fig. 4b and Extended Data Fig. 7a,b). Functional analysis further shows that LolA recruitment must remain dynamic. Disrupting the LolF hook strongly impairs lipoprotein transfer despite substantial residual ATPase activity (Fig. 3f and Extended Data Fig. 7e), whereas the F94E variant remains persistently associated with *Ab*LolDF but is likewise defective in transfer (Fig. 3f,g). Productive handover requires a correctly organized interaction that can form for substrate reception and dissociate during turnover. The *Ab*LolA Δ68-76 phenotype adds a geometric requirement: detectable transporter association is retained, but transfer is strongly impaired (Fig. 3f and Fig. 3g).

The abaucin-bound structures allow comparison with two recent structural studies of *Ab*LolDF. Zhang et al. resolved two- and four-abaucin complexes and proposed that sequential inhibitor binding drives stepwise opening of the transmembrane cavity to generate an induced-fit inhibitory pocket. Pang et al. instead emphasized a two-ligand state in which two abaucin molecules occupy symmetry-related sites and trap a substrate-accessible but transport-incompetent transporter. We observe a wide cavity accommodating four abaucin molecules and a narrower state retaining two (Fig. 5a,b). Analysis of the wide-state particles revealed an additional level of ligand heterogeneity. In the C1 reconstruction of the four-ligand state, the two central abaucin molecules are not equivalent and are compatible with distinct folded and extended poses. Abaucin 3 is also less well resolved than the other ligands, with density that does not fully encompass its trifluoromethyl group, indicating greater conformational uncertainty at this end of the molecule. Reprocessing the same particles with C2 symmetry makes the two central sites equivalent and produces similar intermediate ligand conformations (Extended Data Fig. 10b,c). The twofold protein architecture therefore does not require equivalent ligand poses. Symmetry averaging can obscure asymmetric inhibitor configurations.

This comparison suggests that the different abaucin configurations reported across studies need not represent mutually exclusive binding modes. They may represent different members of a conformational ensemble or different symmetry-averaged views. Cavity shape, ligand occupancy and reconstruction symmetry can each affect the apparent poses. In our wide state, the two peripheral abaucin molecules occupy sites retained in the two-ligand narrow state, whereas the additional central ligands are less tightly constrained (Fig. 5b-d and Extended Data Fig. 10b,c). The weaker density associated with the trifluoromethyl-containing end of abaucin 3 is consistent with greater conformational heterogeneity at these central sites. We therefore propose that the peripheral sites form a relatively stable inhibitory core, while the central sites become populated in more expanded cavity conformations and tolerate multiple ligand poses. This arrangement offers a structural connection between the two- and four-ligand states (Fig. 5d).

The narrow abaucin-bound state combines substrate-like global cavity dimensions with an inhibitor-specific door-bar geometry. Its lower cavity also differs from LolA-engaged state 2 (Figs. 4b and 5d), while its A377 separation is larger than that of uninhibited *Ab*LolDF-*Ab*LolB (Extended Data Fig. 10d). Functional observations support the same interpretation. Abaucin inhibits lipoprotein transfer from proteoliposomes reconstituted with preassembled *Ab*LolDF-BamE complexes (Fig. 5f), binds *Ab*LolDF and *Ab*LolDF-*Ab*LolB with similar apparent affinities (Extended Data Fig. 10e), and loses potency against the A377T and Y378A variants (Fig. 5e,f). Because BamE was already associated with LolDF before abaucin exposure in the transfer assay, inhibition does not require prevention of initial substrate association. These findings are consistent with abaucin engaging a substrate-associated transporter and diverting it into a non-productive conformation that cannot complete handover to LolA.

LolDF and LolCDE organize functional asymmetry through different architectures. LolDF generates functional asymmetry dynamically within a homodimer (Fig. 2b and Fig. 3c-e), whereas LolCDE encodes a related division of labour through two divergent transmembrane subunits^14,15,23^. These differences also create distinct druggable conformations. Lolamicin preferentially targets LolCDE-containing bacteria, whereas abaucin exploits the repeated surfaces and conformational plasticity of *Ab*LolDF^16,19^. In this framework, substrate-induced asymmetry organizes normal lipoprotein handover, while abaucin redirects the same conformational flexibility into non-productive wide and narrow states (Fig. 5a-d and Extended Data Fig. 10f). These differences identify distinct structural contexts for selective inhibitors of lipoprotein transport.

## Methods

### Protein expression and purification

Genes encoding LolD and LolF from *Acinetobacter baumannii* ATCC 19606, together with the indicated variants, were synthesized by GenScript and cloned into pET28a with a C-terminal His_6_ tag on LolD. Recombinant proteins were expressed in *Escherichia coli* C43 cells. Cultures were grown to an OD_600_ of approximately 0.6, induced with 150 μM IPTG and incubated for 16 h at 18 °C with shaking at 120 r.p.m. Cells were harvested and resuspended in 500 mM NaCl, 20 mM HEPES, pH 8.0, and 20 mM imidazole. Cells were disrupted by high-pressure homogenization in the presence of 1% (v/v) Protease Inhibitor Cocktail (TargetMol). Membranes were collected by ultracentrifugation at 200,000 × g for 1 h at 4 °C and solubilized for 2 h at room temperature in the same buffer supplemented with 1% (w/v) n-dodecyl-β-D-maltopyranoside (DDM; Anatrace). Insoluble material was removed by centrifugation at 50,000 × g for 1 h at 4 °C. The supernatant was loaded onto Ni-NTA resin (Smart-Lifesciences), washed with 500 mM NaCl, 20 mM HEPES, pH 8.0, 50 mM imidazole and 0.01% (w/v) lauryl maltose neopentyl glycol (LMNG), and eluted with the same buffer containing 300 mM imidazole. Wild-type and mutant *Ab*LolDF complexes were further purified by size-exclusion chromatography on a Superdex 200 Increase 10/300 GL column (Cytiva) equilibrated in 200 mM NaCl, 20 mM HEPES, pH 8.0, and 0.005% (w/v) LMNG. Peak fractions were pooled and concentrated for subsequent biochemical or cryo-EM experiments.

*Ab*LolDF-BamE and *Ab*LolDF-LolB complexes, together with the indicated variants, were produced by co-expression in *E. coli* C43 cells from pRSFDuet constructs encoding His_6_-tagged LolDF and Strep-tagged lipoprotein cargo. The complexes were purified using the same membrane preparation, detergent extraction, Ni-NTA affinity chromatography and size-exclusion chromatography procedures described above for *Ab*LolDF.

His_6_-tagged *Ab*LolA(29-231) and His_6_-GST-*Ab*LolA(29-231) constructs, together with the indicated variants, were cloned into pET28a and expressed in *E. coli* C43 cells. Cultures were grown to an OD_600_ of approximately 0.6, induced with 150 μM IPTG and incubated for 16 h at 18 °C with shaking at 120 r.p.m. Cells were harvested and resuspended in 500 mM NaCl, 20 mM HEPES, pH 8.0, and 20 mM imidazole. Cells were disrupted by high-pressure homogenization, and insoluble material was removed by centrifugation at 50,000 × g for 1 h at 4 °C. The clarified lysate was loaded onto Ni-NTA resin (Smart-Lifesciences). The resin was washed with 500 mM NaCl, 20 mM HEPES, pH 8.0, and 50 mM imidazole, and bound proteins were eluted with the same buffer containing 300 mM imidazole. His_6_-*Ab*LolA and His_6_-GST-*Ab*LolA proteins were further purified by size-exclusion chromatography on a Superdex 200 Increase 10/300 GL column equilibrated in 200 mM NaCl and 20 mM HEPES, pH 8.0. Peak fractions were pooled and concentrated for subsequent biochemical or structural experiments.

### Site-directed mutagenesis

Point mutations and deletions in *Ab*LolDF and *Ab*LolA were introduced using the ClonExpress II One Step Cloning Kit (Vazyme). Plasmids were isolated using the PurePlasmid Mini Kit (CW0500M, CWBIO), and all constructs were verified by Nanopore Sequencing (CWBIO) before protein expression and purification.

### In-gel digestion and LC-MS/MS

SEC-purified nucleotide-free *Ab*LolDF was separated by SDS-PAGE, and the entire protein-containing region of the gel lane was excised. The excised gel was processed, and proteins were digested in gel with trypsin. The resulting in-gel-digested peptides were identified by liquid chromatography-tandem mass spectrometry, performed by Bio-Tech Pack Technology. Spectra were searched against *Ab*LolDF and *E. coli* protein databases. *E. coli* lipoproteins were assigned using UniProtKB lipoprotein annotations^24^. MS1 area and unique-peptide counts were used to summarize protein identifications. MS1 area was treated as a semi-quantitative signal and was not converted to molar abundance.

### Purification of the *Ab*LolDF-*Ab*LolB-*Ab*LolA ternary complex

Purified LolDF-LolB was incubated with GST-LolA(29-231) at a molar ratio of 1:5 for 1 h at 4°C. The resulting ternary complex was isolated by GST-affinity chromatography and further purified on a Superdex 200 Increase 10/300 column. Fractions containing LolDF-LolB-LolA were pooled and concentrated to 3 mg/mL.

### Abaucin

Abaucin (RS102895; MedChemExpress, cat. no. HY-18611A; CAS no. 300815-41-2) was purchased from MedChemExpress and used for biochemical, structural and growth-inhibition experiments.

### ATPase measurements

ATP hydrolysis by *Ab*LolDF was measured using a malachite-green phosphate assay. Unless otherwise indicated, reactions contained 200 mM NaCl, 20 mM HEPES, pH 7.5, 5 mM MgCl_2_, 0.01% (w/v) LMNG and 0.16 mg/mL *Ab*LolDF. Reactions were initiated by addition of ATP and incubated for 30 min at 37 °C. ATP hydrolysis was terminated by addition of an equal volume of 0.5 M EDTA. Malachite-green reagent was then added, and samples were incubated for 30 min at 25 °C before absorbance was measured at 630 nm.

For ATP-dependent activity measurements, wild-type *Ab*LolDF and the indicated LolD catalytic variants were assayed over the indicated range of ATP concentrations. For analysis of the *Ab*LolDF^Δhook^ variant, ATPase activity was measured under the same reaction conditions. For abaucin inhibition experiments, reactions contained 2 mM ATP and the indicated concentrations of abaucin. ATPase activities at each abaucin concentration were normalized to the corresponding activity measured in the absence of abaucin, and concentration-response curves were fitted in Prism 10 (GraphPad) to determine IC_50_ values. Each condition was measured in three technical replicates in each of three independent experiments.

### Proteoliposome reconstitution and lipoprotein-transfer assay

Proteoliposomes containing *Ab*LolDF-BamE were prepared using *Escherichia coli* polar lipids (Avanti Polar Lipids). Purified *Ab*LolDF-BamE complexes were used directly for reconstitution, such that the BamE cargo was already associated with *Ab*LolDF before initiation of the transfer assay. Lipids dissolved in chloroform were dried by rotary evaporation and further dried under vacuum overnight. The lipid film was hydrated to 10 mg/mL in buffer A containing 200 mM NaCl and 20 mM HEPES, pH 7.5. The lipid suspension was sonicated and extruded through a 0.2 μm polycarbonate membrane to generate unilamellar liposomes. Liposomes were destabilized with LMNG and incubated with purified wild-type or mutant *Ab*LolDF-BamE for 1 h on ice. The detergent concentration was reduced by 150-fold dilution with buffer A, and the resulting proteoliposomes were collected by ultracentrifugation at 300,000 × g for 30 min at 4 °C. Proteoliposome pellets were resuspended in buffer A to a final lipid concentration of 1 mg/mL.

For lipoprotein-transfer assays, proteoliposomes reconstituted with preassembled *Ab*LolDF-BamE complexes were mixed with purified GST-tagged wild-type *Ab*LolA(29-231) or the indicated *Ab*LolA variants at an *Ab*LolDF:*Ab*LolA molar ratio of 1:21. Where indicated, reactions contained 2 mM ATP and 5 mM MgCl_2_, 5 mM EDTA, or 0.4 mM abaucin. Reactions were incubated for 20 min at 25 °C and terminated by addition of 7.5 volumes of ice-cold buffer A. Proteoliposomes were pelleted by ultracentrifugation at 300,000 × g for 30 min at 4 °C, and the GST-*Ab*LolA-containing supernatant was collected. *Ab*LolA-associated BamE was subsequently isolated by GST-affinity pull-down. Fractions were analysed by SDS-PAGE and immunoblotting using antibodies against GST and the Strep-tagged lipoprotein cargo. Signals were quantified using a Tanon-5200 Multi imaging system. Unless otherwise indicated, assays were performed in three independent experiments, each with three technical replicates.

### Proteoliposome co-sedimentation assay

Association of *Ab*LolA with membrane-reconstituted *Ab*LolDF was assessed by proteoliposome co-sedimentation. *Ab*LolDF proteoliposomes were prepared as described above and adjusted to 1 mg/mL in buffer A containing 200 mM NaCl and 20 mM HEPES, pH 7.5. Proteoliposomes were incubated with purified His_6_-GST-tagged wild-type *Ab*LolA(29-231) or the indicated *Ab*LolA variants (F94E, W96E, D117A, D227A and Δ68-76) at an *Ab*LolDF:*Ab*LolA molar ratio of 1:21. Where indicated, proteoliposomes containing *Ab*LolDF^Δhook^ were used in place of wild-type *Ab*LolDF.

Following incubation, reactions were diluted 7.5-fold with ice-cold buffer A and centrifuged at 300,000 × g for 30 min at 4 °C. The supernatant containing soluble *Ab*LolA was collected, and the proteoliposome pellet was washed once with ice-cold buffer A and centrifuged again under the same conditions. The final pellet was resuspended in a volume of buffer A equal to that of the corresponding supernatant. Equivalent proportions of the pellet and supernatant fractions were analysed by SDS-PAGE and immunoblotting using antibodies against GST-tagged *Ab*LolA. *Ab*LolA signals in the pellet and supernatant fractions were quantified using a Tanon-5200 Multi imaging system. Each assay was performed in three independent experiments.

### Abaucin growth-inhibition assay

The growth-inhibitory activity of abaucin against *Acinetobacter baumannii* ATCC 19606 was assessed in liquid culture. Bacteria were streaked on LB agar and incubated overnight at 37 °C. Three individual colonies were used to inoculate 5 ml LB cultures, which were grown overnight at 37 °C with shaking at 220 r.p.m. Overnight cultures were diluted 1:100 into fresh LB and grown for approximately 3 h at 37 °C until the OD_600_ reached 0.4-0.6. Cultures were then diluted 1:100 into fresh LB, and 200 μl aliquots were transferred to 96-well plates. Abaucin was added at concentrations ranging from 0.5 to 16 μg/mL. OD_600_ was recorded hourly for 12 h using a Spark multimode microplate reader (Tecan) with shaking at 37 °C. Wells containing medium only were used for background subtraction. For the analysis shown in Fig. 5g, OD600 values at 4 h were used. Each biological replicate was measured in technical triplicate, and data were analysed using Prism 10 (GraphPad).

### SPR analysis

Surface plasmon resonance (SPR) measurements were performed using a Biacore 1K instrument (GE Healthcare). Purified *Ab*LolDF or *Ab*LolDF-*Ab*LolB was immobilized on CM5 sensor chips, and abaucin was injected as a serial dilution over a concentration range of 0.14-300 μM. Sensorgrams were processed and analysed using Biacore Insight software. Apparent equilibrium dissociation constants (K_d_) were obtained by steady-state affinity analysis. Experiments were performed independently three times, and the reported Kd values represent the mean of the three independent measurements.

### Cryo-EM sample preparation and data collection

For the nucleotide-free sample, LolDF was adjusted to 8.6 mg/mL. An aliquot of 3.5 μL was applied to a glow-discharged Quantifoil holey gold grid (R 1.2/1.3, Au 300 mesh). The grid was glow-discharged at 25 mA for 60 s. Grid preparation was performed at 16 °C and 100% humidity using a Vitrobot Mark IV (Thermo Fisher Scientific). A wait time of 15 s was used before the sample was blotted for 4.5 s at blot force 4. The grid was plunge-frozen in liquid ethane cooled by liquid nitrogen.

For the AMP-PNP-bound LolDF sample, purified LolDF was supplemented with 2 mM AMP-PNP and 5 mM MgCl_2_. An aliquot of 3.5 μL at 3.75 mg/mL was applied to a glow-discharged Quantifoil holey gold grid (R 1.2/1.3, Au 300 mesh). The grid was glow-discharged at 25 mA for 60 s and prepared at 16 °C and 100% humidity using a Vitrobot Mark IV. A wait time of 15 s was used before the sample was blotted for 3 s at blot force 4. The grid was plunge-frozen in liquid ethane cooled by liquid nitrogen.

For the ATP-bound LolD^E172Q^F sample, purified protein was supplemented with 2 mM ATP and 5 mM MgCl_2_. An aliquot of 3.5 μL at 4 mg/mL was applied to a glow-discharged Quantifoil holey gold grid (R 1.2/1.3, Au 300 mesh). The grid was glow-discharged at 25 mA for 60 s and prepared at 16 °C and 100% humidity using a Vitrobot Mark IV. A wait time of 30 s was used before the sample was blotted for 3 s at blot force 4. The grid was plunge-frozen in liquid ethane cooled by liquid nitrogen.

For the LolDF-LolB sample, purified complex was supplemented with 2 mM ATP and 1 mM MgCl_2_. An aliquot of 3.5 μL at 15 mg/mL was applied to a glow-discharged Quantifoil holey gold grid (R 1.2/1.3, Au 300 mesh). The grid was glow-discharged at 25 mA for 60 s and prepared at 16 °C and 100% humidity using a Vitrobot Mark IV. A wait time of 30 s was used before the sample was blotted for 3 s at blot force 2. The grid was plunge-frozen in liquid ethane cooled by liquid nitrogen.

For the LolDF-LolB-LolA sample, purified complex was supplemented with 2 mM AMP-PNP and 5 mM MgCl_2_. An aliquot of 3.5 μL at 2 mg/mL was applied to a glow-discharged Quantifoil holey gold grid (R 1.2/1.3, Au 300 mesh). The grid was glow-discharged at 25 mA for 60 s and prepared at 16 °C and 100% humidity using a Vitrobot Mark IV. A wait time of 15 s was used before the sample was blotted for 3 s at blot force 2. The grid was plunge-frozen in liquid ethane cooled by liquid nitrogen.

For the narrow abaucin-bound dataset, purified LolDF-LolB was supplemented with 2 mM ATP, 5 mM MgCl_2_ and 1 mM abaucin. An aliquot of 3.5 μL at 8 mg/mL was applied to a glow-discharged Quantifoil holey gold grid (R 1.2/1.3, Au 300 mesh). The grid was glow-discharged at 25 mA for 60 s and prepared at 16 °C and 100% humidity using a Vitrobot Mark IV. A wait time of 30 s was used before the sample was blotted for 3 s at blot force 2. The grid was plunge-frozen in liquid ethane cooled by liquid nitrogen.

For the LolDF-abaucin dataset that yielded the predominant wide class and a minor narrow class, LolDF purified without intentional lipoprotein co-expression was supplemented with 2 mM ATP, 5 mM MgCl_2_ and 1 mM abaucin. An aliquot of 3.5 μL at 7 mg/mL was applied to a glow-discharged Quantifoil holey gold grid (R 1.2/1.3, Au 300 mesh). The grid was glow-discharged at 25 mA for 60 s and prepared at 16 °C and 100% humidity using a Vitrobot Mark IV. A wait time of 30 s was used before the sample was blotted for 2 s at blot force 2. The grid was plunge-frozen in liquid ethane cooled by liquid nitrogen.

Cryo-EM data were collected on a Titan Krios G3i transmission electron microscope (Thermo Fisher Scientific) operated at 300 kV. The microscope was equipped with a Falcon 4i direct electron detector, and data acquisition was controlled using EPU software (Thermo Fisher Scientific). All movies were recorded in counting mode. The nominal magnification, calibrated pixel size, electron dose, defocus range, exposure time and number of frames for each dataset are provided in Supplementary Data Table 1.

### Cryo-EM image processing, model building and structural analysis

Motion correction and contrast-transfer-function (CTF) estimation were performed in cryoSPARC v4.3.0^25^. Micrographs with estimated CTF resolution worse than 6 Å were excluded during manual exposure curation. Two-dimensional classification, three-dimensional classification and refinement were also performed in cryoSPARC. The two half-datasets were refined independently. Global resolution was estimated at the gold-standard Fourier shell correlation (FSC) threshold of 0.143^26^. Local resolution was calculated in cryoSPARC and visualized in UCSF ChimeraX^27^.

For nucleotide-free LolDF, 7,078 movies were collected and 6,784 micrographs were retained after motion correction and CTF estimation. A total of 756,625 particles were extracted, and the final reconstruction reached 3.4 Å (Extended Data Fig. 2). For LolD^E172Q^F-ATP, 681 movies yielded 668 retained micrographs and 78,337 extracted particles; the final reconstruction reached 3.6 Å. For LolDF-AMP-PNP, 6,986 movies yielded 6,906 retained micrographs and 524,791 extracted particles; the final reconstruction reached 2.8 Å (Extended Data Fig. 3).

For LolDF-LolB-ATP, 7,389 movies yielded 6,849 retained micrographs and 739,634 extracted particles. The final reconstruction reached 3.1 Å (Extended Data Fig. 4). For LolDF-LolB-LolA, 5,516 movies yielded 5,400 retained micrographs and 659,511 extracted particles. Further classification produced two reconstructions at 2.9 Å and 2.7 Å (Extended Data Fig. 6).

For LolDF-abaucin purified without intentional lipoprotein co-expression, 7,114 movies yielded 6,924 retained micrographs and 403,400 extracted particles. Three-dimensional classification produced a predominant wide class and a minor narrow class. The predominant wide class was refined to 3.6 Å (Extended Data Fig. 9a-c). The same wide-state particle set was additionally reprocessed with C2 symmetry to assess the effect of symmetry enforcement on ligand-density interpretation; this reconstruction was used only as a processing control. For the LolDF-LolB-abaucin sample, 8,147 movies yielded 7,866 retained micrographs and 413,513 extracted particles. Duplicate removal retained 171,609 particles, which produced a 3.4 Å narrow reconstruction (Extended Data Fig. 9d-f).

Initial models were predicted using AlphaFold 3^28^. The highest-ranked model from five predictions was selected for each complex and fitted into the corresponding cryo-EM density map by rigid-body placement in UCSF ChimeraX^27^. Models were subsequently rebuilt manually in Coot to match the experimental density and the sequences of the constructs used^29^. Nucleotides and other ligands were modelled according to the corresponding map densities. Atomic models were refined against the cryo-EM maps using phenix.real_space_refine with appropriate stereochemical and secondary-structure restraints^30^. Reference-model restraints were used during model optimization where appropriate but were not applied uniformly to all final refinements. For the abaucin-bound structures, ligand-specific geometry restraints were applied during refinement. Model geometry was assessed using MolProbity within Phenix^31^.

## Lead contact

Further information and requests for resources and reagents should be directed to and will be fulfilled by the lead contact, Zhaoxing Li.

## Materials availability

Plasmids and constructs generated in this study are available from the corresponding authors upon reasonable request.

## Data and code availability

Atomic coordinates for the structures reported in this study have been deposited in the Protein Data Bank under accession codes 45GN (nucleotide-free AbLolDF), 45GO (AbLolDE172QF– ATP), 45GP (AbLolDF–AMP-PNP), 45GQ (AbLolDF–AbLolB–ATP), 45GR (AbLolDF–AbLolB–GST–AbLolA state 1), 45GS (AbLolDF–AbLolB–GST–AbLolA state 2), 45GT (AbLolDF–abaucin, wide-cavity state) and 45GU (AbLolDF–AbLolB–abaucin, narrow-cavity state). The corresponding cryo-EM maps have been deposited in the Electron Microscopy Data Bank under accession codes EMD-83252, EMD-83253, EMD-83254, EMD-83255, EMD-83256, EMD-83257, EMD-83258 and EMD-83259, respectively. Source data are provided with this paper.

These data are publicly available as of the date of publication. This paper does not report original code.

Any additional information required to reanalyse the data reported in this paper is available from the lead contact upon request.

## ACKNOWLEDGEMENTS

This work is supported by Innovative Drug Research and Development-National Science and Technology Major Project (2025ZD1802300, Y.X.), National Natural Science Foundation of China (82625061, 82473977, Y.X.; 82304614, Z.Li), the Natural Science Foundation of Jiangsu Province (BK20240094, Y.X.; BK20260204, M.L.), Postgraduate Research & Practice Innovation Program of Jiangsu Province (26CXJH1746, J.K.) and National Innovation and Entrepreneurship Training Program for Undergraduate (S202610316010, Z.Lin). We would like to thank the Instrument Analysis Center (IAC) at Shanghai Jiao Tong University and Center of Cryo-Electron Microscopy at Nanjing Medical University for Cryo-EM data collection.

## AUTHOR CONTRIBUTIONS

Conceptualization: Z.X.L., Y.B.X., J.P.K.

Funding acquisition: Y.B.X., Z.X.L., M.L.L., J.P.K., Z.L.

Investigation: J.P.K., Z.Y.Z., Z.L., N.Y., R.Y.H.

Methodology: J.P.K., Y.B.X., Z.X.L., H.F.S., Z.Y.Z., H.Y., Z.L.

Project administration: Y.B.X., J.P.K., Z.X.L., H.F.S.

Resources: Y.B.X., Z.X.L., H.F.S., M.L.L., M.R.C.

Supervision: Z.X.L., Y.B.X., J.P.K., H.F.S., H.F.G., M.R.C.

Validation: J.P.K., Z.X.L., Z.Y.Z., Z.L., N.Y.

Visualization: J.P.K., Z.X.L., Z.L., Z.Y.Z.

Writing-original draft: J.P.K., Z.X.L., Y.B.X., Z.Y.Z., Z.L.

Writing-review & editing: Z.X.L., Y.B.X., J.P.K.

## Competing interests

Authors declare that they have no competing interests.

## Notes

### Competing Interest Statement

The authors have declared no competing interest.

