## Supplemental figure for "Substrate-induced asymmetry coordinates lipoprotein handover and abaucin trapping in *Acinetobacter baumannii* LolDF"

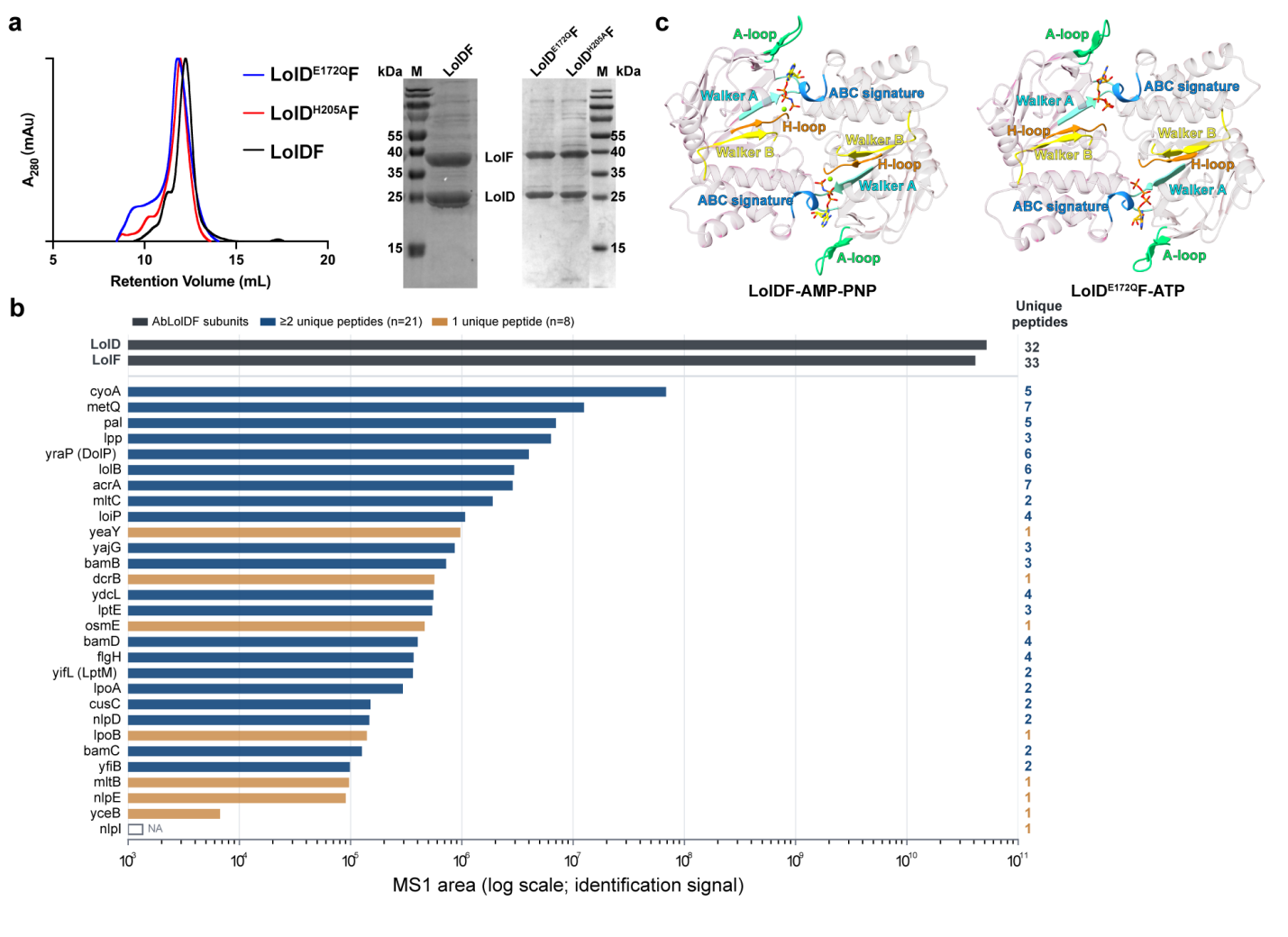
**Extended Data Fig. 1 | Purification and proteomic characterization of *Ab*LolDF.**

**a,** Size-exclusion chromatography profiles and SDS-PAGE analysis of wild-type *Ab*LolDF and the *Ab*LolDE172QF and *Ab*LolDH205AF variants.

**b,** LC-MS/MS analysis of SEC-purified nucleotide-free *Ab*LolDF. Bars show MS1 areas for *Ab*LolD, *Ab*LolF and 29 lower-abundance *Escherichia coli* lipoproteins. The number of unique peptides supporting each protein identification is shown on the right. MS1 area is shown as a semi-quantitative identification signal and was not converted to molar abundance.

**c,** Top views of the LolD dimer in *Ab*LolDF-AMP-PNP and *Ab*LolDE172QF-ATP, showing the two nucleotide-binding sites.

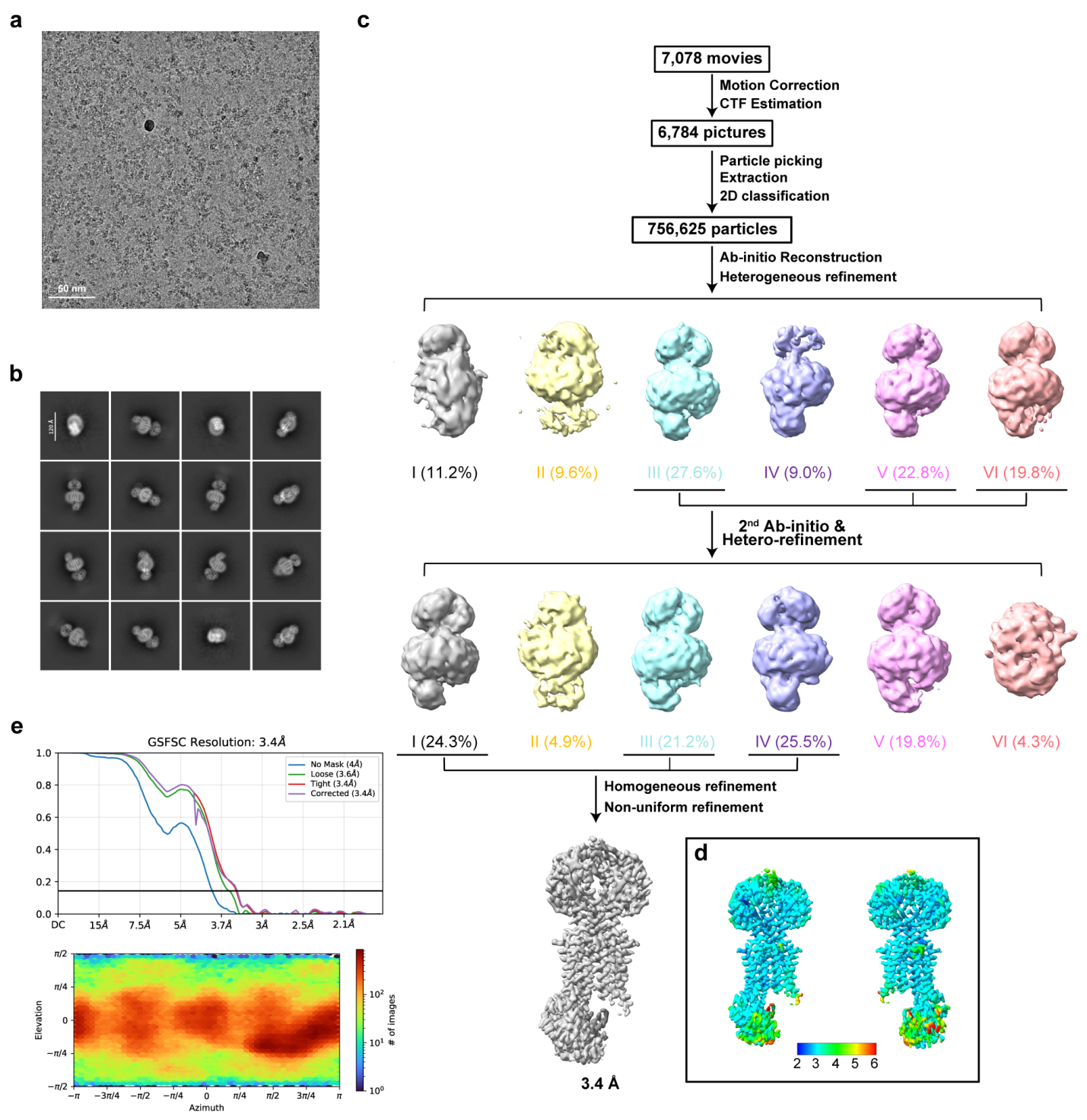

**Extended Data Fig. 2 | Cryo-EM data processing for nucleotide-free *Ab*LolDF.**

**a,** Representative cryo-EM micrograph of nucleotide-free *Ab*LolDF.

**b,** Representative two-dimensional class averages.

**c,** Cryo-EM data-processing workflow.

**d,** Final reconstruction coloured by local resolution.

**e,** Gold-standard Fourier shell correlation curves and particle-orientation distribution for the final reconstruction.

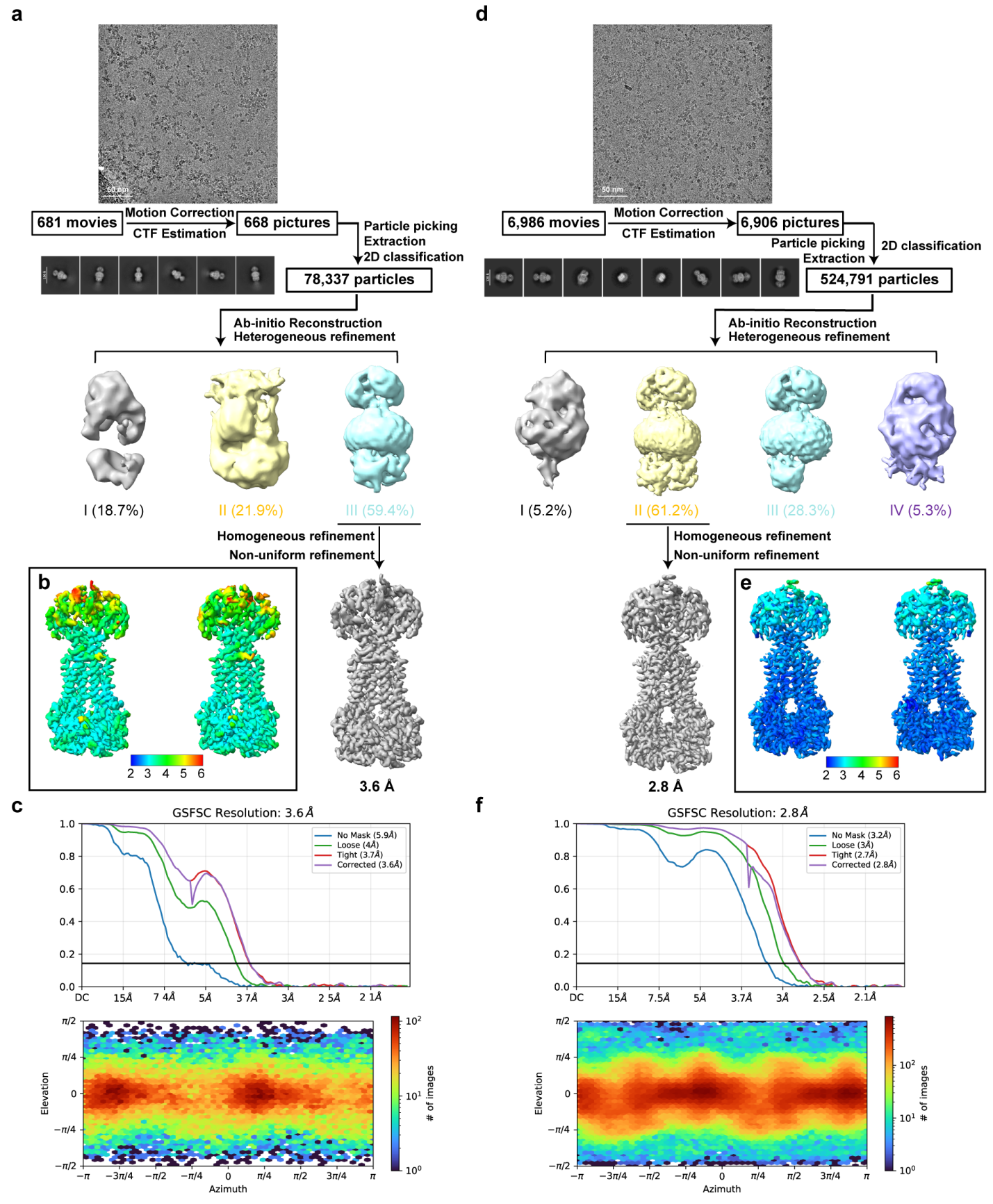

**Extended Data Fig. 3 | Cryo-EM data processing for *Ab*LolDE172QF-ATP and *Ab*LolDF-AMP-PNP.**

**a,** Cryo-EM data-processing workflow for *Ab*LolDE172QF-ATP.

**b,** Final *Ab*LolDE172QF-ATP reconstruction coloured by local resolution.

**c,** Gold-standard Fourier shell correlation curves and particle-orientation distribution for *Ab*LolDE172QF-ATP.

**d,** Cryo-EM data-processing workflow for *Ab*LolDF-AMP-PNP.

**e,** Final *Ab*LolDF-AMP-PNP reconstruction coloured by local resolution.

**f,** Gold-standard Fourier shell correlation curves and particle-orientation distribution for *Ab*LolDF-AMP-PNP.

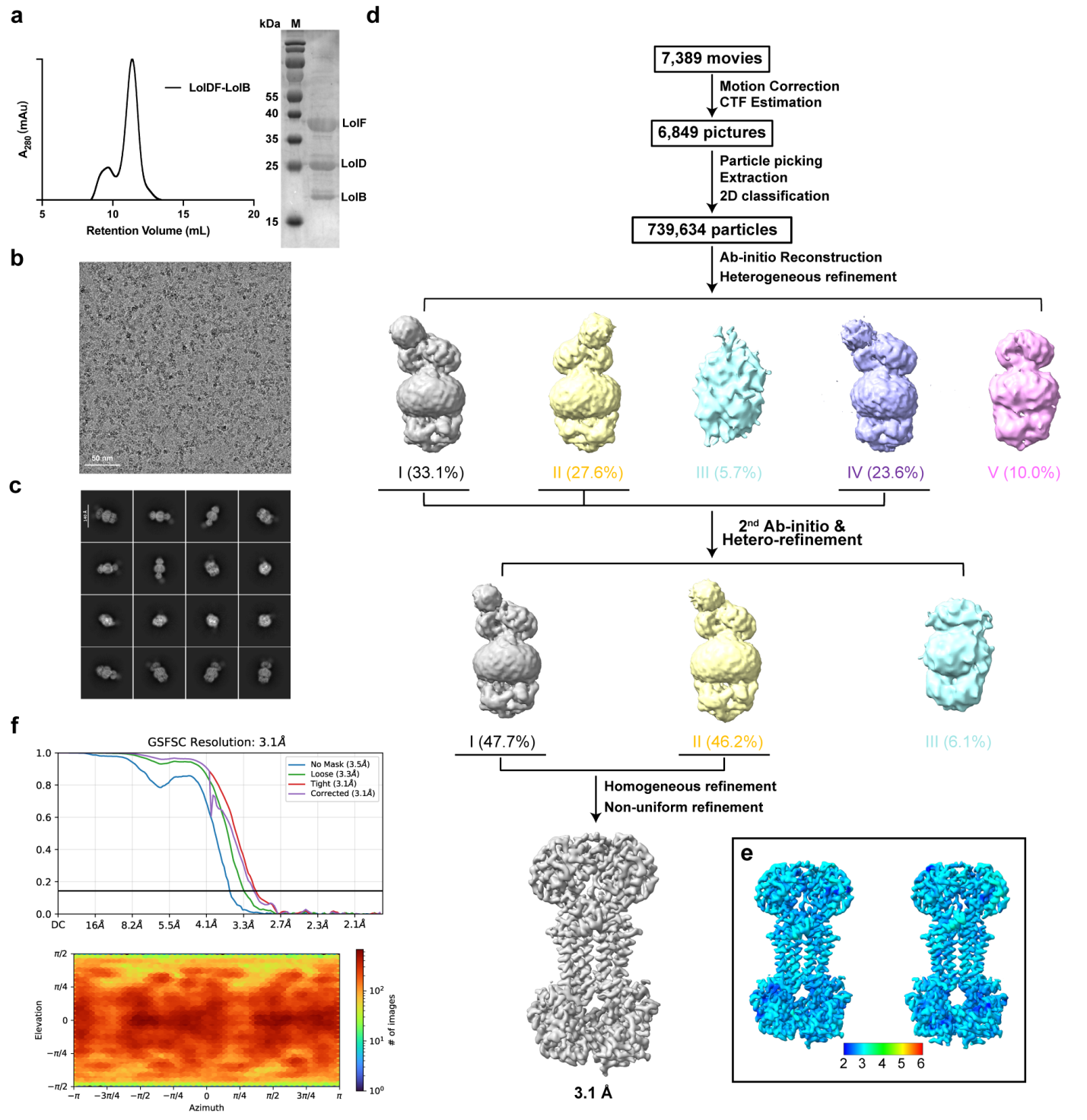

**Extended Data Fig. 4 | Cryo-EM data processing for *Ab*LolDF-*Ab*LolB-ATP.**

**a,** Size-exclusion chromatography profile and SDS-PAGE analysis of the *Ab*LolDF-*Ab*LolB complex.

**b,** Representative cryo-EM micrograph.

**c,** Representative two-dimensional class averages.

**d,** Cryo-EM data-processing workflow.

**e,** Final reconstruction coloured by local resolution.

**f,** Gold-standard Fourier shell correlation curves and particle-orientation distribution for the final reconstruction.

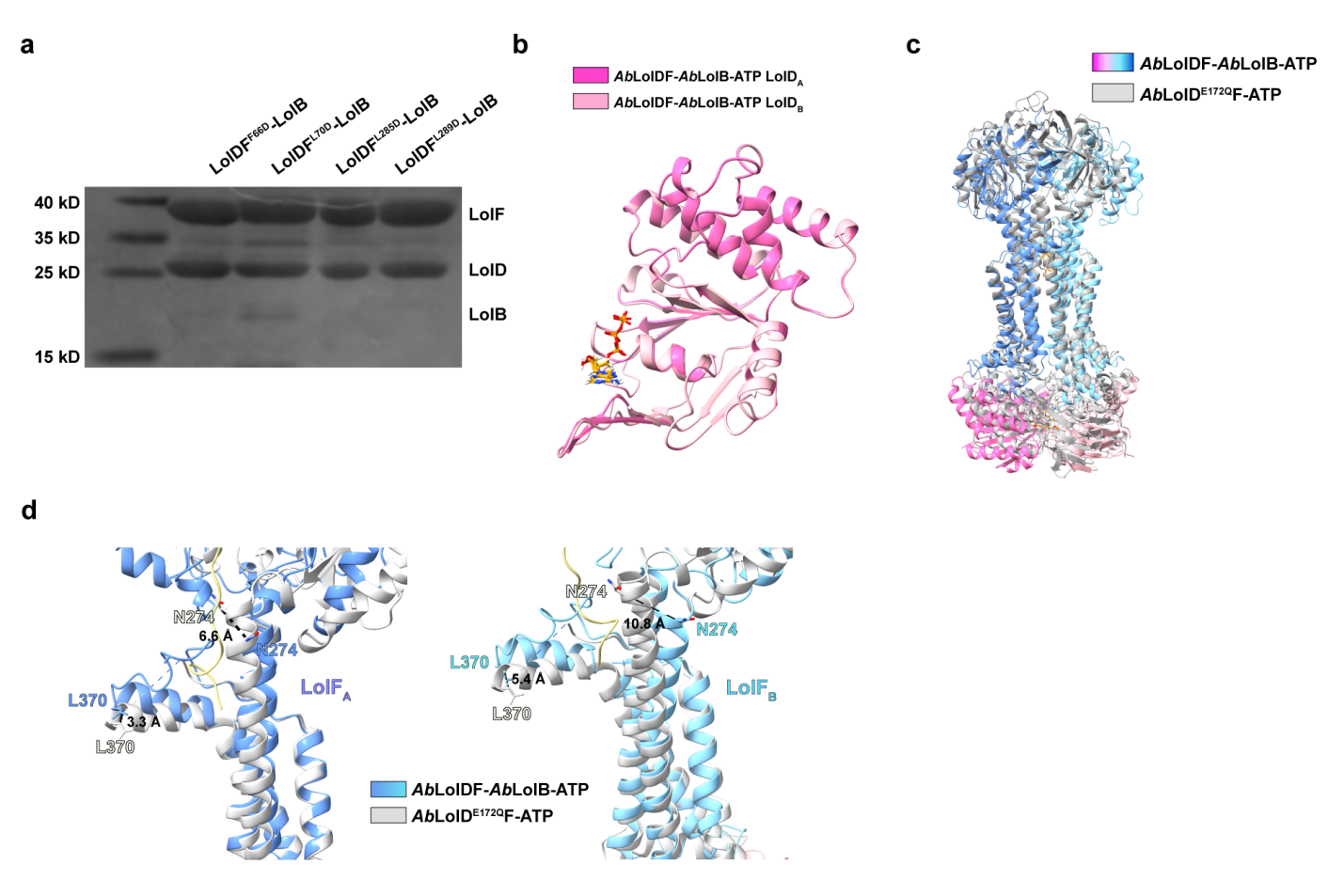
**Extended Data Fig. 5 | Conformational changes associated with *Ab*LolB binding to *Ab*LolDF.**

**a,** Co-purification analysis of the indicated lipoprotein-binding-site variants of *Ab*LolDF.

**b,** Superposition of the two LolD subunits in *Ab*LolDF-*Ab*LolB-ATP.

**c,** Structural comparison of *Ab*LolDF-*Ab*LolB-ATP and *Ab*LolDE172QF-ATP.

**d,** Local comparison of the LolF transmembrane helices in *Ab*LolDF-*Ab*LolB-ATP and *Ab*LolDE172QF-ATP. Selected inter-residue distances are indicated.

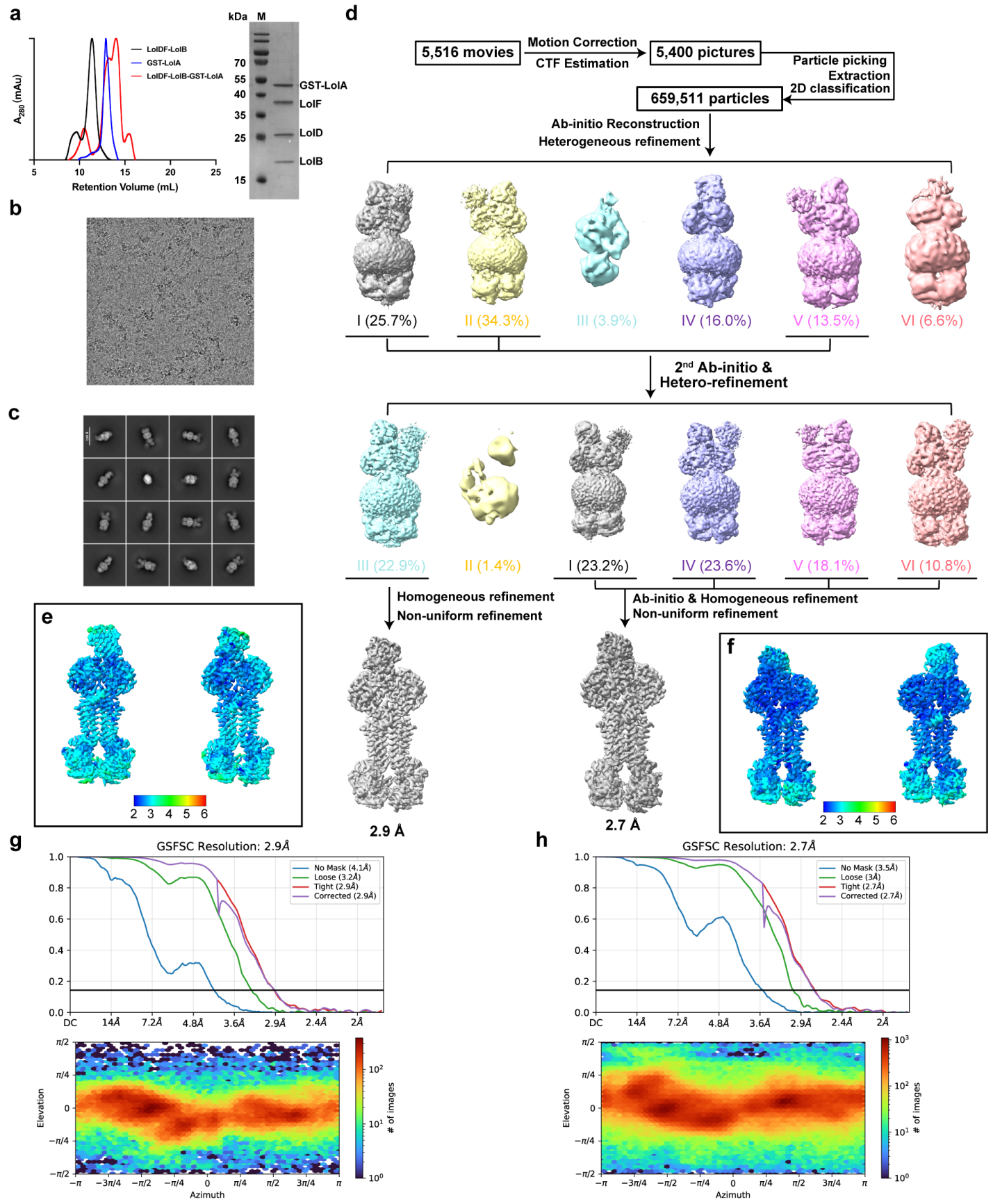
**Extended Data Fig. 6 | Cryo-EM processing of *Ab*LolDF-*Ab*LolB-GST-*Ab*LolA.**

**a,** SEC profiles and SDS-PAGE of the ternary complex and its components.

**b,** Representative cryo-EM micrograph.

**c,** Representative two-dimensional class averages.

**d,** Data-processing workflow yielding two states.

**e,** f, States 1 and 2 coloured by local resolution.

**g,** h, Gold-standard FSC curves and particle-orientation distributions for states 1 and 2.

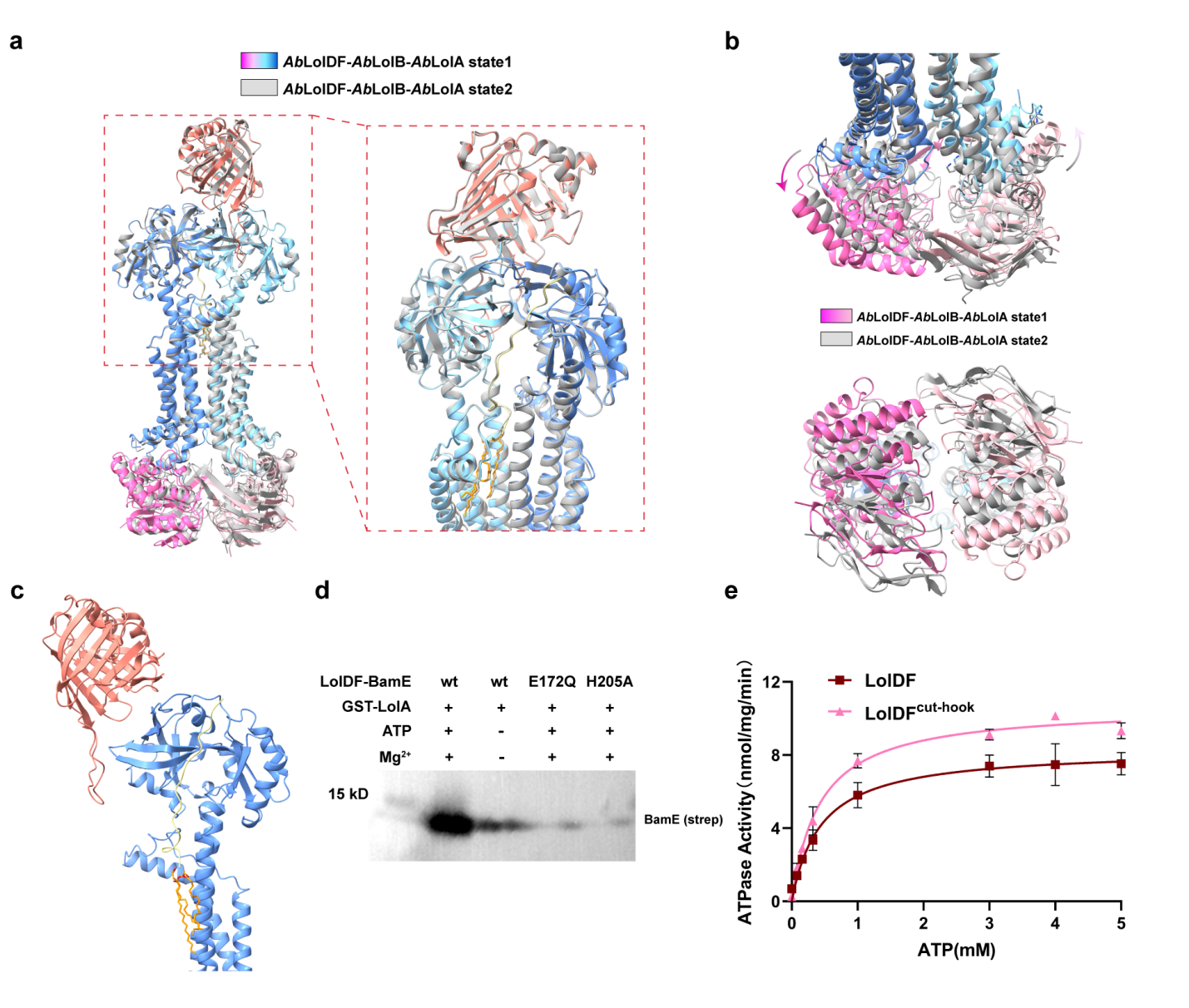
**Extended Data Fig. 7 | Structural comparison of the two *Ab*LolDF-*Ab*LolB-GST-*Ab*LolA states.**

**a,** Structural superposition of states 1 and 2, with an enlarged view of the *Ab*LolA- and *Ab*LolB-binding region.

**b,** Comparison of the LolD dimers in states 1 and 2.

**c,** Overall view of LolF protomer A and its interactions with *Ab*LolA and *Ab*LolB.

**d,** Representative immunoblot showing in vitro lipoprotein transfer mediated by wild-type *Ab*LolDF and the *Ab*LolDE172QF and *Ab*LolDH205AF variants under the indicated ATP and Mg²⁺ conditions. A representative result from three independent experiments is shown; each experiment was performed in technical triplicate, with similar results.

**e,** ATP-dependent ATPase activities of wild-type *Ab*LolDF and *Ab*LolDFΔhook. Data are presented as mean ± s.d. from three independent experiments, each performed in technical triplicate.

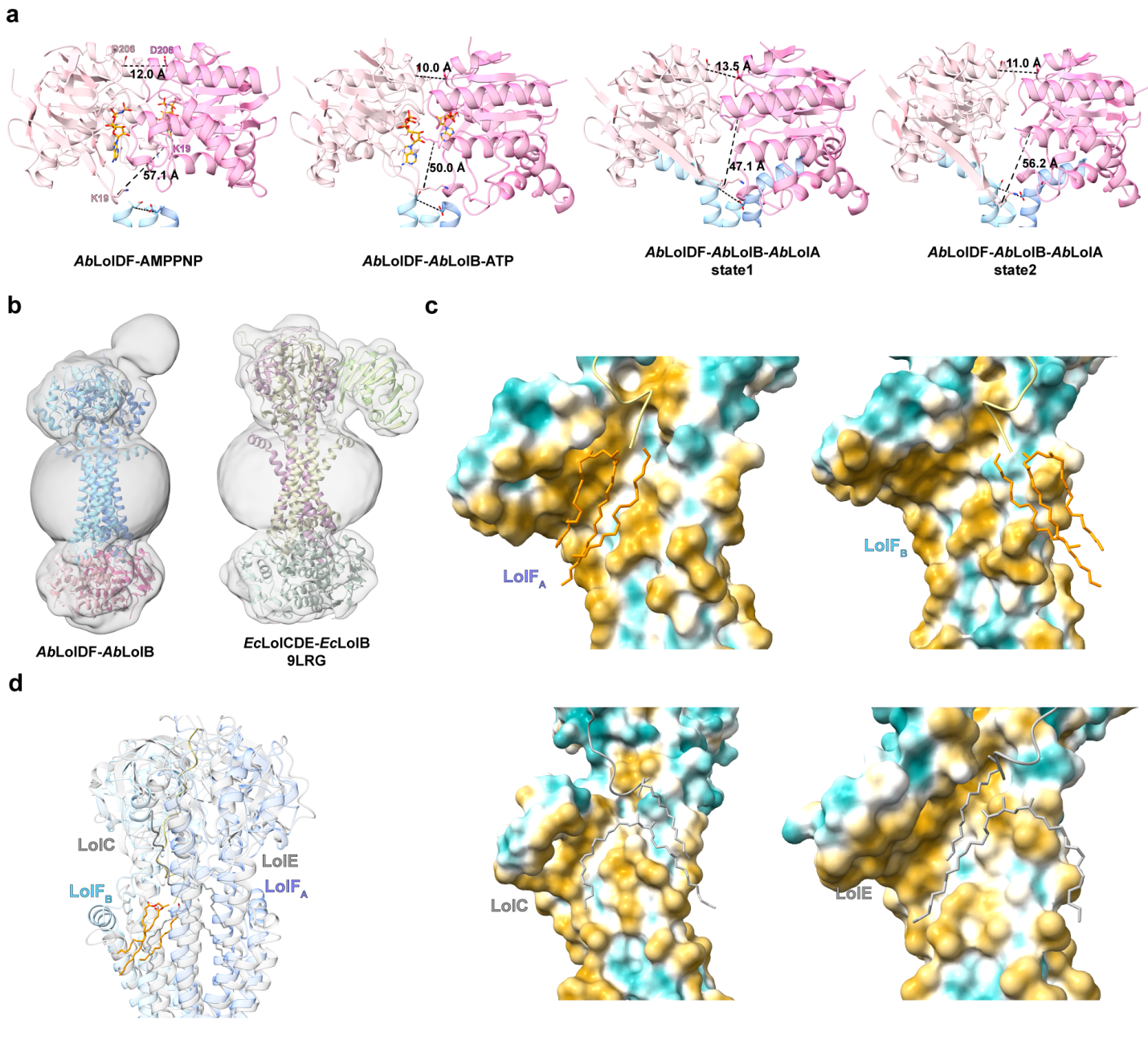
**Extended Data Fig. 8 | Structural comparison of *Ab*LolDF with *Escherichia coli* LolCDE.**

**a,** LolD dimer arrangements in the indicated *Ab*LolDF states. Distances between representative residues quantify changes at the upper and lower regions of the dimer.

**b,** Cryo-EM density map and cartoon representation of the *Ab*LolDF-*Ab*LolB complex and *Escherichia coli* LolCDE-LolB complex (PDB 9RLG).

**c,** Cross-sectional hydrophobicity surfaces of the LolF protomers in *Ab*LolDF-*Ab*LolB and of LolC and LolE in the *Escherichia coli* LolCDE-LolB complex.

**d,** Structural comparison of the transmembrane regions of *Ab*LolDF-*Ab*LolB and *Escherichia coli* LolCDE-LolB.

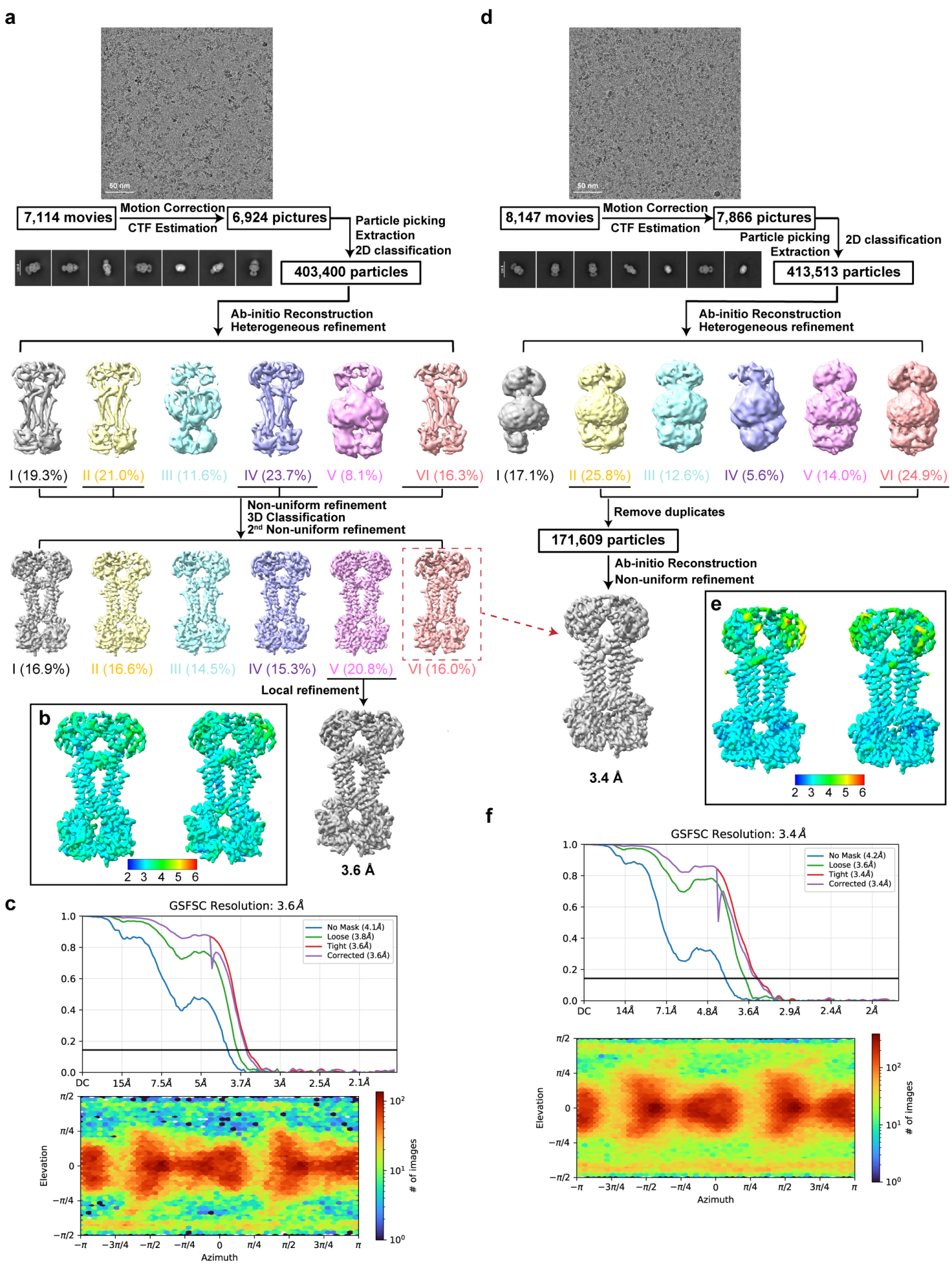
**Extended Data Fig. 9 | Cryo-EM processing of abaucin-bound *Ab*LolDF.**

**a-c,** Data-processing workflow, local-resolution map and gold-standard FSC curves with particle-orientation distribution for the wide-cavity state from the *Ab*LolDF-abaucin sample.

**d-f,** Corresponding analyses of the narrow-cavity state from the *Ab*LolDF-*Ab*LolB-abaucin sample.

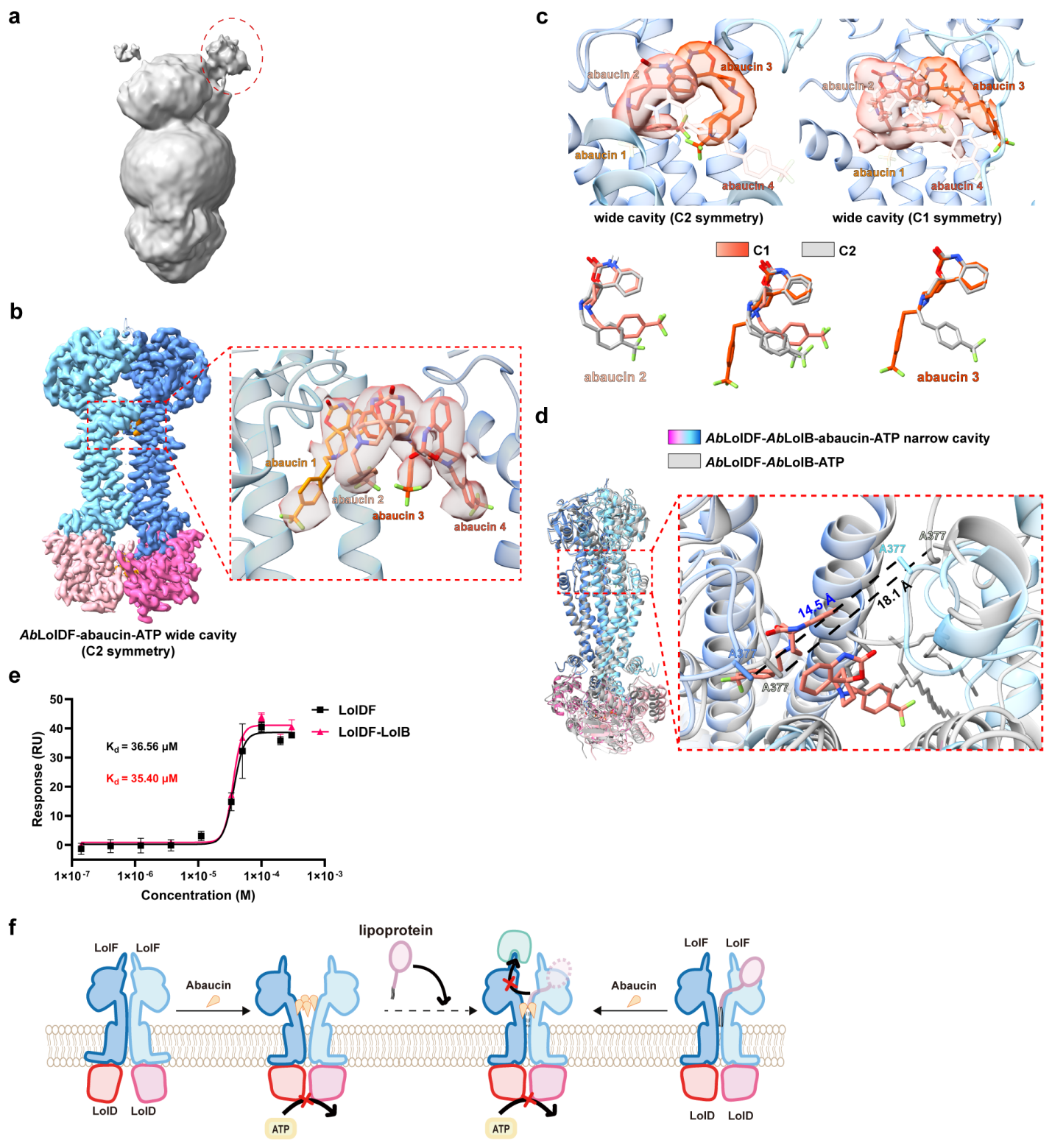
**Extended Data Fig. 10 | Additional analyses of abaucin-bound *Ab*LolDF.**

**a,** Gaussian-smoothed density of the *Ab*LolDF-*Ab*LolB-abaucin reconstruction displayed in UCSF ChimeraX (width = 2.0). The red circle indicates putative lipoprotein-associated density.

**b,** C2-symmetrized reprocessing of the same wide-state particle set used for the final C1 reconstruction, shown as a symmetry-control analysis with an enlarged view of the central abaucin-binding region.

**c,** Comparison of the central abaucin densities and illustrative ligand fits in the final C1 reconstruction and the C2-symmetrized processing control.

**d,** Structural comparison of narrow-cavity *Ab*LolDF-*Ab*LolB-abaucin with *Ab*LolDF-*Ab*LolB-ATP.

**e,** Steady-state surface plasmon resonance (SPR) analysis of abaucin binding to *Ab*LolDF and *Ab*LolDF-*Ab*LolB. Equilibrium responses were fitted by steady-state affinity analysis, and the resulting apparent dissociation constants (Kd) are indicated.

**f,** Schematic models illustrating abaucin inhibition of substrate-poor and putative lipoprotein-associated *Ab*LolDF.
