## Supplemental table for "Substrate-induced asymmetry coordinates lipoprotein handover and abaucin trapping in *Acinetobacter baumannii* LolDF"

**Supplementary Data Table 1**

**Cryo-EM data collection, refinement and validation statistics**

| **Parameter** | **AbLolDF** **nucleotide-free** | **AbLolDF** **AMP-PNP** | **AbLolD(E172Q)F** **ATP** | **AbLolDF–AbLolB** **ATP** |
| --- | --- | --- | --- | --- |
| PDB | 45GN | 45GP | 45GO | 45GQ |
| EMDB | EMD-83252 | EMD-83254 | EMD-83253 | EMD-83255 |
| **Data collection and processing** | | | | |
| Microscope | Titan Krios G3i | Titan Krios G3i | Titan Krios G3i | Titan Krios G3i |
| Detector | Falcon 4i | Falcon 4i | Falcon 4i | Falcon 4i |
| Acquisition software | EPU | EPU | EPU | EPU |
| Magnification | 130k | 130k | 130k | 130k |
| Voltage (kV) | 300 | 300 | 300 | 300 |
| Electron exposure (e^-^/Å^2^) | 50 | 50 | 50 | 50 |
| Defocus range (μm) | 0.8-2.4 | 0.8-2.4 | 0.8-2.4 | 0.6-2.4 |
| Pixel size (Å) | 0.932 | 0.932 | 0.929 | 0.932 |
| Movies collected (no.) | 7,078 | 6,986 | 681 | 7,389 |
| Micrographs retained (no.) | 6,784 | 6,906 | 668 | 6,849 |
| Symmetry imposed | C1 | C1 | C1 | C1 |
| Initial particle images (no.) | 756,625 | 524,791 | 78,337 | 739,634 |
| Final particle images (no.) | 377,592 | 321,119 | 46,538 | 585,790 |
| Map resolution (Å) | 3.4 | 2.8 | 3.6 | 3.1 |
| FSC threshold | 0.143 | 0.143 | 0.143 | 0.143 |
| **Refinement** | | | | |
| Initial model used | AlphaFold 3 | AlphaFold 3 | AlphaFold 3 | AlphaFold 3 |
| Model resolution (Å) | 3.7 | 3.2 | 4.0 | 3.3 |
| FSC threshold | 0.5 | 0.5 | 0.5 | 0.5 |
| Map sharpening B factor (Å^2^) | 40 | 0 | 0 | 0 |
| **Model composition** | | | | |
| Non-hydrogen atoms | 7,675 | 9,729 | 9,715 | 9,933 |
| Protein residues | 1,005 | 1,259 | 1,256 | 1,274 |
| Ligands | 0 | 4 | 2 | 4 |
| **B factors (mean, Å^2^)** | | | | |
| Protein | 56.46 | 74.63 | 74.34 | 120.19 |
| Ligand | n.a. | 73.18 | 62.00 | 119.32 |
| **R.m.s. deviations** | | | | |
| Bond lengths (Å) | 0.004 | 0.004 | 0.004 | 0.003 |
| Bond angles (°) | 0.754 | 0.661 | 0.670 | 0.565 |
| **Validation** | | | | |
| MolProbity score | 2.06 | 2.36 | 2.57 | 1.85 |
| Clashscore | 14.35 | 10.73 | 18.20 | 5.96 |
| Poor rotamers (%) | 0.83 | 3.22 | 2.56 | 1.50 |
| **Ramachandran plot** | | | | |
| Favored (%) | 93.88 | 93.58 | 91.64 | 94.29 |
| Allowed (%) | 6.02 | 6.26 | 8.04 | 5.24 |
| Disallowed (%) | 0.10 | 0.16 | 0.32 | 0.48 |

| **Parameter** | **AbLolDF–AbLolB** **GST–AbLolA** **state 1** | **AbLolDF–AbLolB** **GST–AbLolA** **state 2** | **AbLolDF–abaucin** **wide** | **AbLolDF–AbLolB** **abaucin narrow** |
| --- | --- | --- | --- | --- |
| PDB | 45GR | 45GS | 45GT | 45GU |
| EMDB | EMD-83256 | EMD-83257 | EMD-83258 | EMD-83259 |
| **Data collection and processing** | | | | |
| Microscope | Titan Krios G3i | Titan Krios G3i | Titan Krios G3i | Titan Krios G3i |
| Detector | Falcon 4i | Falcon 4i | Falcon 4i | Falcon 4i |
| Acquisition software | EPU | EPU | EPU | EPU |
| Magnification | 130k | 130k | 130k | 130k |
| Voltage (kV) | 300 | 300 | 300 | 300 |
| Electron exposure (e^-^/Å^2^) | 50 | 50 | 50 | 50 |
| Defocus range (μm) | 0.6-2.4 | 0.6-2.4 | 0.8-2.2 | 1.0-2.2 |
| Pixel size (Å) | 0.932 | 0.932 | 0.932 | 0.929 |
| Movies collected (no.) | 5,516 | 5,516 | 7,114 | 8,147 |
| Micrographs retained (no.) | 5,400 | 5,400 | 6,924 | 7,866 |
| Symmetry imposed | C1 | C1 | C1 | C1 |
| Initial particle images (no.) | 659,511 | 659,511 | 403,400 | 413,513 |
| Final particle images (no.) | 111,265 | 366,841 | 67,245 | 171,609 |
| Map resolution (Å) | 2.9 | 2.7 | 3.6 | 3.4 |
| FSC threshold | 0.143 | 0.143 | 0.143 | 0.143 |
| **Refinement** | | | | |
| Initial model used | AlphaFold 3 | AlphaFold 3 | AlphaFold 3 | AlphaFold 3 |
| Model resolution (Å) | 3.1 | 2.9 | 3.7 | 3.5 |
| FSC threshold | 0.5 | 0.5 | 0.5 | 0.5 |
| Map sharpening B factor (Å^2^) | 0 | 0 | 0 | 0 |
| **Model composition** | | | | |
| Non-hydrogen atoms | 11,480 | 11,464 | 9,936 | 9,789 |
| Protein residues | 1,484 | 1,482 | 1,270 | 1,256 |
| Ligands | 2 | 2 | 6 | 4 |
| **B factors (mean, Å^2^)** | | | | |
| Protein | 117.25 | 129.98 | 66.45 | 130.42 |
| Ligand | 108.22 | 126.91 | 61.72 | 130.74 |
| **R.m.s. deviations** | | | | |
| Bond lengths (Å) | 0.005 | 0.004 | 0.003 | 0.003 |
| Bond angles (°) | 0.867 | 0.637 | 0.594 | 0.631 |
| **Validation** | | | | |
| MolProbity score | 2.22 | 2.20 | 1.89 | 1.83 |
| Clashscore | 14.20 | 11.24 | 8.92 | 7.26 |
| Poor rotamers (%) | 1.60 | 1.92 | 1.13 | 0.24 |
| **Ramachandran plot** | | | | |
| Favored (%) | 93.82 | 93.61 | 94.53 | 93.41 |
| Allowed (%) | 5.71 | 5.99 | 5.31 | 6.35 |
| Disallowed (%) | 0.48 | 0.41 | 0.16 | 0.24 |
